# DNA gyrase in live bacteria forms liquid condensates through weak multivalent bonding of excess GyrB

**DOI:** 10.64898/2026.08.13.744598

**Authors:** Aisha H Syeda, Katy Hollands, Lewis Frame, Jack Shepherd, Alex Payne-Dwyer, Eleanor Goffee, Nicolas Burton, Aakash Basu, Agnes Noy, Anthony Maxwell, Mark C Leake

**Affiliations:** School of Physics, Engineering and Technology, University of York, York, YO10 5DD, UK; Bioscience Technology Facility, Department of Biology, University of York, York, YO10 5DD, UK; Department of Biosciences, South Road, Durham, DH1 3LE, UK; Inspiralis Limited, Innovation Centre, Norwich Research Park, Colney Lane, Norwich, NR4 7GJ, UK; Department of Molecular Microbiology, John Innes Centre, Norwich Research Park, Norwich, NR4 7UH, UK; Department of Biology, University of York, York, YO10 5DD, UK

## Abstract

Type IIA bacterial topoisomerase DNA gyrase, a GyrA/GyrB heterotetramer, has crucial roles maintaining transcription and DNA replication by relaxing positive DNA supercoils through introducing negative supercoils. However, rates of gyrase-catalysed supercoiling *in vitro* cannot explain much higher rates required *in vivo*. To address this puzzle, we used high-speed single-molecule fluorescence imaging of GyrA/GyrB reporters in live *Escherichia coli*, indicating that cells contain ∼40% more GyrB than GyrA expressed in a diffuse pool or in clusters whose mobility depends on whether they are bound to DNA. Unexpectedly, we discovered that clusters are non-stoichiometric containing ∼150% more GyrB than GyrA, significantly greater than the cellular average, with fluorescence recovery after photobleaching revealing that clustered GyrA and GyrB behave as a liquid whose abundance can be increased by applying gyrase-targeting antibiotics. Structural docking indicates that the liquid state is stabilised through excess GyrB progressively binding to existing clusters via weak, multivalent interactions. By operating in liquid condensates, A_2_B_2_ that dissociates from DNA can rebind rapidly instead of diffusing away, increasing enzyme processivity to enable multiple rounds of catalysis that can keep pace with transcription and DNA replication *in vivo*. This demonstrates a new role for condensates of overcoming kinetic limitations imposed by diffusion.

**Graphical abstract:** 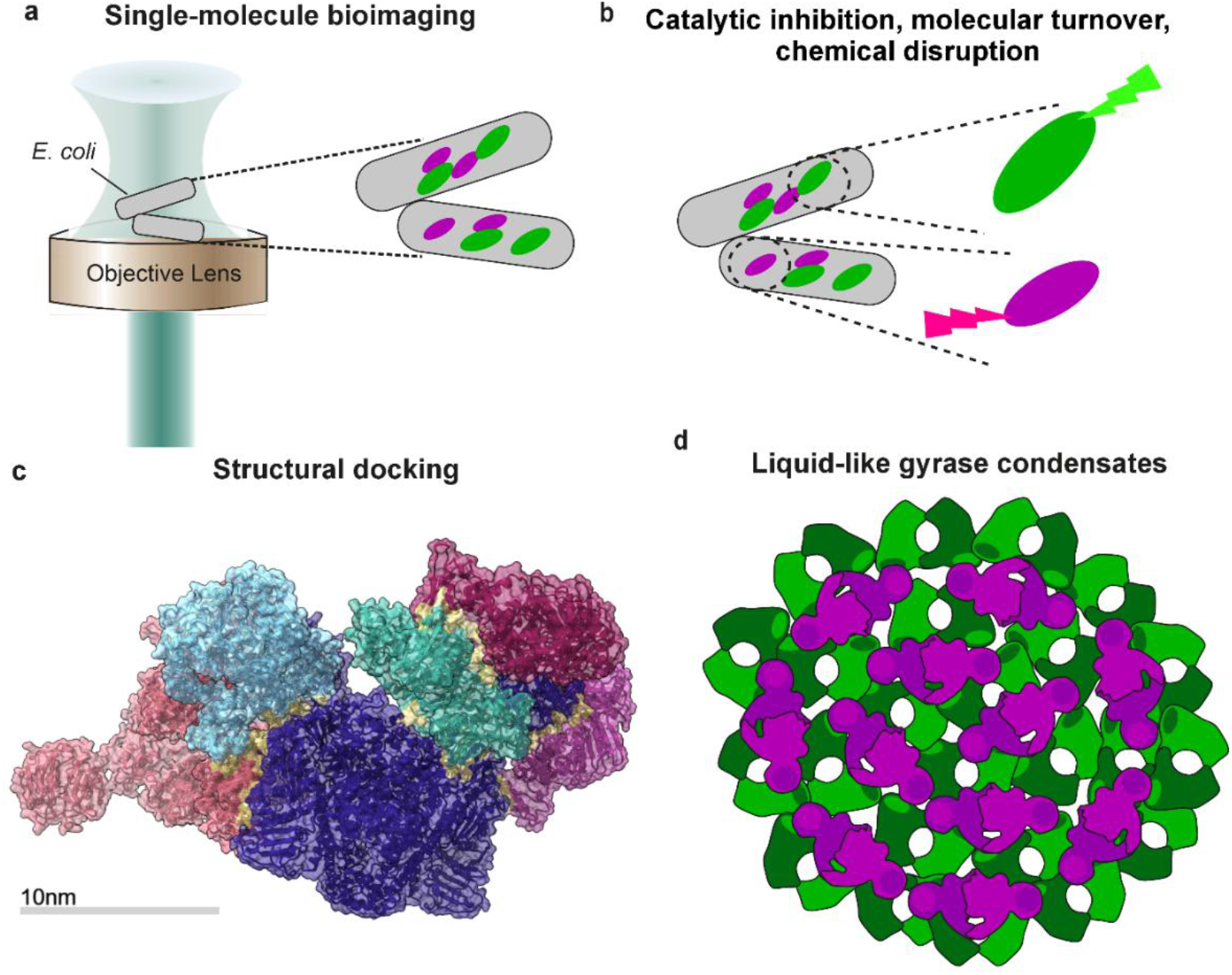

## Introduction

In bacteria, progression of DNA replication and transcription generates positively (+) supercoiled DNA ahead of either DNA polymerase or RNA polymerase respectively^1^ which constitutes a physical barrier to these crucial cellular processes. In bacteria, this is resolved by two essential type II topoisomerases, DNA topoisomerase IV, believed to act primarily to decatenate entangled DNA, and DNA gyrase^2^, which changes DNA topology by introducing (−) negative supercoils^2,3^. Extensive structural and biochemical data *in vitro* indicate that gyrase’s catalytically active unit is a complex comprising two GyrA and two GyrB subunits forming a heterotetramer (A_2_B_2_) which acts via a strand-passage mechanism^4^ in which it introduces a double-strand (ds) break into a DNA segment bound at the GyrA dimer interface resulting in a transient GyrA-DNA cleavage complex^2^. Then, the enzyme passes another DNA segment through this ds break, driven by the binding and hydrolysis of ATP. Following conclusion of the catalytic cycle with religation of the ds break, this process results in an overall decrease in DNA linking number by 2.

Gyrase is unique in this ability to introduce (−) supercoils into DNA and is thus a central enzyme for maintaining supercoiling homeostasis in bacteria^5–8^. Homeostasis maintains a net (−) supercoiled state for the *E. coli* chromosome, with this level important for regulation of most DNA processes including transcription, repair/recombination and replication^9,10^; the expression of several genes, including gyrase, is regulated through supercoiling^11^. However, gyrase activities must be responsive to these processes taking place in different spatial locations of the chromosome since local supercoiling constantly undergoes alteration by ongoing transcription, repair/recombination and replication^12,13^.

Transcription generates (+) supercoiling of DNA in front of translocating RNA polymerase ^1,14,15^. Despite the higher abundance of RNA polymerase in *E. coli*, DNA replication likely generates more severe topological barriers, as the bidirectional translocation of two replisomes around the circular chromosome produces (+) supercoils far more rapidly (∼1,000 base pairs per second bp/s^16,17^) than transcription (∼60 bp/s^7,8,12^). DNA (+) supercoiling ahead of the DNA polymerase is balanced by (-) supercoiling behind with overall replisome rotation around the DNA helix largely negligible^1^ resulting in accumulation of (+) supercoils ahead of a replication fork. To maintain replication, gyrase needs to relax up to ∼100 (+) supercoils per second for each fork, assuming ∼10 bp per supercoil. However, the catalytic cycle of gyrase has been measured at just ∼1 per second from *in vitro* biochemical assays^18–24^. Since each cycle removes 2 (+) supercoils there may need to be up to ∼50 enzymes acting simultaneously at each fork to keep up with replication *in vivo*. Early studies of antibiotic-mediated *E. coli* chromosome fragmentation^25^ have suggested clustering of gyrase near replication forks broadly consistent with this expectation, while single-molecule magnetic tweezers experiments have demonstrated that *E. coli* gyrase *in vitro* is capable of performing several tens of catalytic events within a single burst prior to dissociating from DNA, albeit with no direct visualisation of bound gyrase and thus not being able to determine if bursts are due to a single bound DNA gyrase A_2_B_2_ heterotetramer or to a cluster^20^. Our earlier study focusing on single-molecule fluorescence imaging of GyrA in live *E. coli*^26^ detected clusters that contained dimeric GyrA over a range of stoichiometry values consistent with a mean of 12 DNA gyrase heterotetramers but extending to as many as ∼50, with the majority of GyrA colocalised with the replisome albeit with no direct visualisation of GyrB. The rate of GyrA dissociation in the vicinity of replisome-colocalised clusters estimated from photoactivated localisation microscopy (PALM) was significantly lower than in other regions of the cell. This observation was consistent with increased enzyme processivity in the vicinity of clusters but offered no direct explanation to its cause.

Here, we use high-speed single-molecule fluorescence microscopy to now visualise both the GyrA and GyrB components simultaneously in the same living cell using monomeric GFP (mGFP) and mCherry genomically-encoded fluorescent-fusion constructs under expression of the native gyrase promoters. We observe GyrA and GyrB to be expressed in either a diffuse cellular pool or in distinct clusters. Stepwise photobleaching analysis of the fluorescent protein tags reveals that the mean B:A ratio overall in cells is 1.4:1 whereas within clusters this ratio is 2.5:1, indicating a significant preferential incorporation of GyrB in clusters. Fluorescence recovery after photobleaching (FRAP) in combination with chemical disruption reveals that GyrA and GyrB within clusters behave as a liquid. The abundance of GyrB can be increased by applying antibiotics that specifically target gyrase activity either through inhibition of ATP binding or stabilisation of the DNA cleavage state. We measure the mobility of GyrA and GyrB within clusters to be dependent on their state of binding to DNA. However, the average GyrB mobility is lower than that of GyrA; this is consistent with structural docking indicating that excess GyrB within clusters forms multiple weak, multivalent links to pre-existing gyrase heterotetramer clusters via neighbouring GyrA. Taken together, our findings offer a new explanation for the high levels of gyrase processivity in living cells, through formation of liquid condensates that are stabilised by weak, multivalent bridges mediated through excess GyrB.

## Results

### *E. coli* cells express 40% more GyrB than GyrA

To study the spatiotemporal dynamics of gyrase *in vivo*, we generated a dual-labelled fluorescent *E. coli* strain comprising GyrA tagged with mCherry and GyrB with monomerised GFP, mGFP. Since both *gyrA* and *gyrB* are essential genes, recovery of viable cells carrying the required fluorescent fusions indicated that the fusions were functional. We find that the generation time of the dual-labelled strain is marginally longer than the wild type (Supplementary Table 1, Supplementary Fig. 1) but well within expectations for being phenotypically functional. The sensitivity to rifampicin, an antibiotic that does not target gyrase, was comparable to wild type; the sensitivity to two gyrase-targeting antibiotics was tested, that to the fluoroquinolone antibiotic ciprofloxacin was comparable to wild type while that to the aminocoumarin coumermycin A1 increased ∼10-fold (Supplementary Fig. 2). The catalytic activity was corroborated by *in vitro* biochemical activity data using the purified fusion constructs to test their ability to generate (−) supercoiled DNA from a relaxed DNA plasmid sample (Supplementary Fig. 3).

Using a single-molecule precise optical imaging technique called Slimfield^27^, we measured fluorescence in the dual-labelled strain corresponding to mCherry and mGFP fluorescent reporters for GyrA and GyrB in all cells (Fig. 1a). Slimfield utilises fluorescence microscopy for detection of single fluorescent proteins with rapid millisecond sampling to track immobile and diffusive fluorescently labelled particles inside living cells^28–31^. In this dual-labelled strain, we quantified the total GyrA and GyrB copy numbers (the number of molecules of each protein per cell) using an approach described previously for a GyrA-mYPet labelled strain^26^ which utilises molecular counting via stepwise photobleaching of single fluorescent proteins^32^. In brief, this method used summation of all cellular fluorescence intensities minus contributions from background noise and autofluorescence background, normalised to the single-molecule brightness of the appropriate fluorescent protein^33^. This analysis indicated a mean of 471 ± 16 (±s.e.m.) GyrA and 656 ± 26 GyrB per cell (Fig. 1b, Table 1). This statistically significant difference indicates that on a cell-by-cell basis *in vivo* there is a ∼40% excess of GyrB molecules compared to GyrA (Student’s t-test, P = 1.30 × 10^−5^). The range of GyrA (∼70-1,380) and GyrB (∼40-2,240) per cell measured from cells cultured in low fluorescence minimal media broadly agreed with previous estimations using electron microscope immunogold labelling of ∼1,000-3,000 molecules per cell using *E. coli* cultured in rich nutrient media^34^. The lower mean levels observed in our study are consistent with reduced DNA replication activity in minimal media requiring less gyrase to resolve detrimental (+) supercoiling in light of known sensitivity of gyrase promoters to DNA supercoiling^12^.

**Figure 1.**
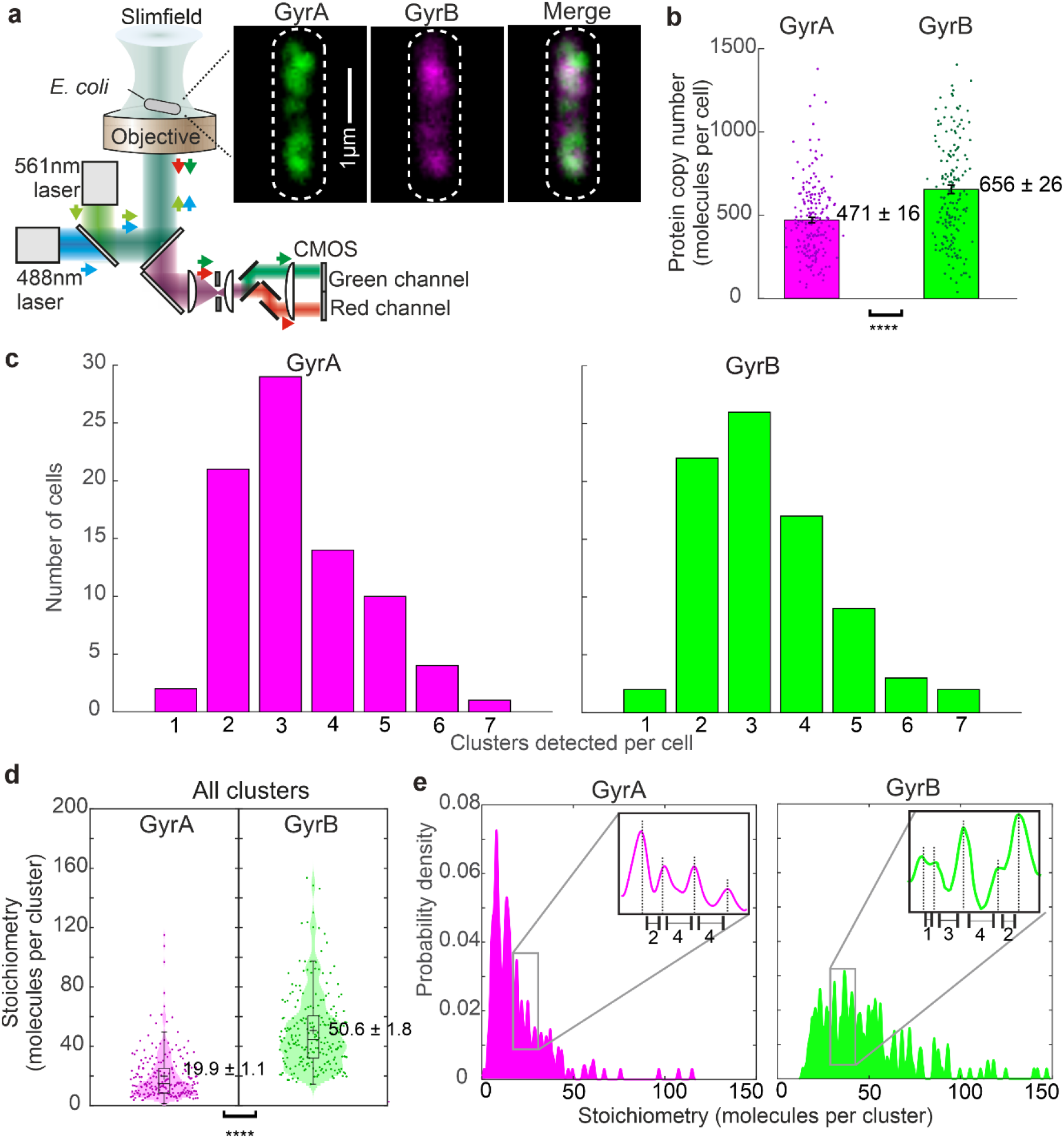
Live *E. coli* bacteria contain significantly more GyrB than GyrA. **a.** Dual colour Slimfield enables single-molecule quantification of GyrA-mCherry (magenta) and mGFP-GyrB (green) in separate colour channels and tracking of gyrase clusters with millisecond sampling; cell outline indicated (white dash), white on merge panel indicates pixel locations in which both colour channels are colocalised. **b.** Distributions for cell copy numbers (number of proteins per cell) overlaid against mean histogram bars for GyrA-mCherry and mGFP-GyrB (difference between GyrA and GyrB is extremely significant, P = 1.3 × 10^−5^), s.e.m. error bars, number of cells n = 81. Here and subsequently, we assume the convention of ns (not significant P≥0.05), * (significant, P<0.05), ** (very significant, P<0.01), *** (highly significant, P<10^−3^), **** (extremely significant, P<10^−4^). **c.** Distribution of number of clusters per cell for GyrA-mCherry (magenta) and mGFP-GyrB (green). Stoichiometry distributions for GyrA and GyrB corresponding to all detected clusters shown as **d.** violin plots overlaid with boxplots (here and subsequently boxes indicate upper and lower quartiles, s.d. error bars, “+” = mean, horizontal bar = median), mean (grey dash line) ±s.e.m. indicated for stoichiometries of all clusters (difference between GyrA and GyrB is extremely significant, P = 1.4 ×10^−38^), and **e.** as kernel density estimations^36^, insets showing zoom-ins with example interval spacings.

**Table 1.**
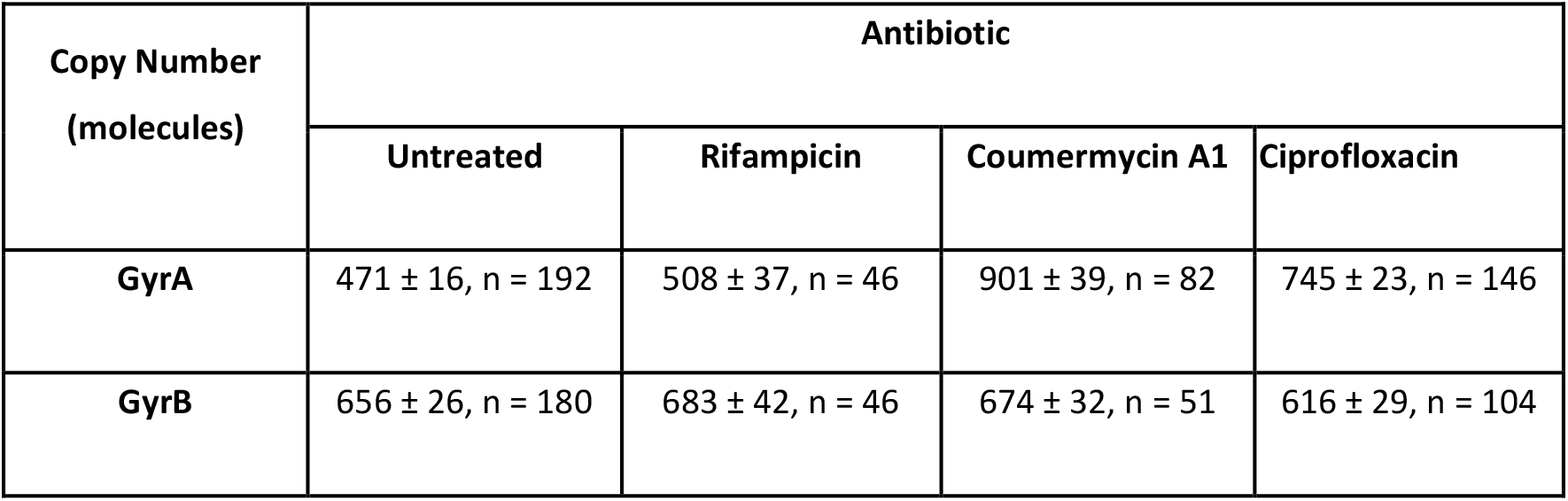
Gyrase subunit protein copy numbers in untreated and antibiotic-treated cells, mean ± s.e.m. error with number n of cells indicated. Treatment with the aminocoumarin coumermycin A1 and fluoroquinolone ciprofloxacin antibiotics significantly increases GyrA copy number (P = 3.76 × 10^−19^ – ciprofloxacin; P = 1.35 × 10^−17^ – coumermycin A1) but does not have a significant impact on GyrB copy number (P = 0.31 – ciprofloxacin; P = 0.66 – coumermycin A1, not significant). Treatment with rifampicin, an antibiotic that does not target gyrase has no significant impact on gyrase subunit copy numbers (P = 0.36 GyrA; P = 0.58 GyrB, not significant).

### Gyrase clusters are significantly enriched with GyrB over GyrA compared to their relative abundance in the whole cell

We observed distinct clusters in each respective colour detection channel (Fig. 1c), the majority of which contained a mixture of GyrA and GyrB, of between 1-7 with a mean 2.3 ± 0.1 clusters per cell for both. Clusters occupied an effective radius of typically a few hundred nm on fluorescence images, comparable to the 300 nm point spread function width of the microscope, with no obvious cellular localisation bias to different regions of the cell. Using in-house software^35^ to automatically track clusters, we found that GyrA and GyrB components of clusters exhibited qualitatively similar heterogeneous spatial localisation to that previously observed for GyrA alone, overlap analysis of the detector channels^36^ indicating that most GyrA and GyrB clusters are colocalised (65 ± 13 % of GyrA clusters are colocalised to a GyrB cluster, 54 ± 11 % of GyrB clusters are colocalised to a GyrA cluster). Our previous observations^26^ with labelled GyrA and replisome marker DnaN revealed that 80% of GyrA colocalises to the replisome. Recent Slimfield imaging of a cell strain co-expressing fluorescently-labelled GyrB and DnaN suggests that 53% of GyrB clusters are colocalised with the replisome^37^. Since in our present study we find that most GyrA and GyrB clusters are colocalised we infer that clusters which contain GyrA and GyrB are likely to be preferentially colocalised to the replisome.

To determine the stoichiometry values of GyrA and GyrB (the number of each respective molecule in a cluster) we used stepwise photobleaching counting applied to individual clusters. We found that GyrA and GyrB have a broad distribution of stoichiometry values (Fig. 1d, e) that does not significantly depend on their state of colocalisation with each other (Supplementary Fig. 4a). Across the whole cluster population, there was a mean of 19.9 ± 1.1 GyrA per cluster (± s.e.m, n = 217) but 50.6 ± 1.8 GyrB per cluster (n = 223) (P = 1.4 × 10^−38^, Supplementary Table 2). The interval between adjacent peaks in the GyrA stoichiometry distribution was consistent with a minimum spacing of approximately two molecules as had been observed in our earlier findings using just a GyrA fluorescent reporter for DNA gyrase^26^, while the stoichiometry distribution pattern for GyrB, although indicating some similar dimeric intervals was less clearly defined, also comprising odd number intervals including single molecule spacing (Fig. 1e insets). The relative abundance within clusters of GyrB is thus approximately ∼150% that of GyrA, equivalent to a B:A relative stoichiometry of 2.5:1 which is nearly twice that of the cell overall, indicating significant enrichment bias that favours GyrB.

Our observation of the formation of GyrB-enriched condensates within cells, together with their potential functional consequences, prompted us to ask whether simply increasing the abundance of GyrB *in vitro* might alter the biochemical properties of the gyrase:DNA complex. To address this, we compared *in vitro* DNA binding by gyrase assembled under canonical conditions (1:1 GyrA:GyrB), with gyrase assembled in presence of excess GyrB (1:5 GyrA:GyrB), using electrophoretic mobility shift assays (EMSA) (Supplementary Fig. 5). Both conditions produced comparable DNA binding under the conditions tested to within ∼5%. This result also suggests that previous in vitro biochemical and single-molecule studies^38,39^ which have used excess GyrB likely do not need to be re-interpreted in the light of possible condensate formation altering biochemical pathways.

### GyrA clusters have a significantly higher mobility compared to GyrB

Based on our previous findings using fluorescently labelled GyrA^26^, the likely state of gyrase cluster binding to DNA is indicated by its mobility – DNA-bound clusters have a relatively low diffusivity, whereas those transiently bound or freely diffusive in the cytosol are significantly more mobile. We measured the mobility of GyrA and GyrB clusters by fitting the initial region of each measured mean square displacement (MSD) obtained from the tracking analysis to a straight line^40^. The mean diffusivity *D* of GyrA clusters was found to be 0.50 ± 0.05 µm^2^/s while that for GyrB was lower by a factor of ∼2 at 0.27 ± 0.04 µm^2^/s (Fig. 2a, P = 4.0 × 10^−4^) which showed similar trends independent of the state of GyrA/GyrB colocalisation (Supplementary Fig. 6a).

**Figure 2.**
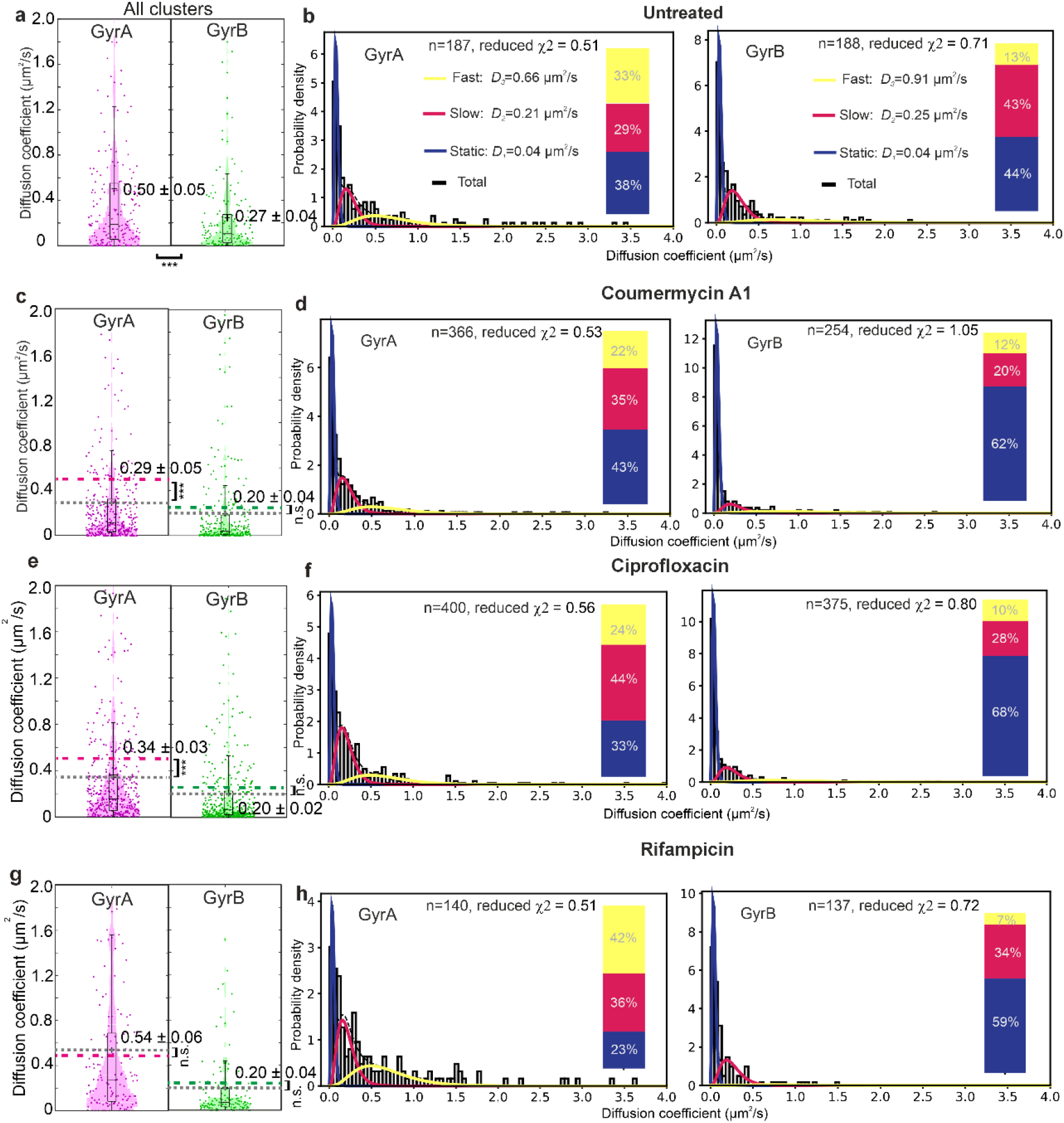
GyrA cluster mobility is higher than GyrB but can be reduced by gyrase-targeting antibiotics. **a.** Violin plots with overlaid jitter data of diffusivity of all clusters for GyrA (magenta, n = 217 clusters) and GyrB (green, n = 223 clusters), mean ± s.e.m. indicated. **b.** Analytical fits to distribution of diffusion coefficients using weighted sum of three diffusion components, modal value of each diffusion coefficient *D_1_*, *D_2_* and *D_3_* and proportion as % of tracks associated with each indicated (inset); reduced χ^2^ value shown for the total fit. Similar analysis for cells in the presence of antibiotics **c,d** coumermycin A1 **e,f** ciprofloxacin; **g,h** rifampicin; mean diffusion coefficient values for untreated data is indicated in magenta and green dashed lines for GyrA and GyrB respectively in panels c., e., g. compared against the associated mean values when treated with each respective antibiotic (grey dashed lines).

We used a biophysical model for the distribution of *D* values^41^ and, as assessed by the reduced χ^2^ and *R*^2^ parameters (Supplementary Fig. 7), were able to fit the data with a weighted sum comprising three diffusion components (Fig. 2b). The “static” diffusion component *D*_1_ had modal values 0.06 and 0.05 μm^2^/s for GyrA and GyrB respectively, consistent with earlier measurements from photoactivated localisation microscopy (PALM) of DNA-bound GyrA^26^; this was comparable with estimates from single-particle tracking of labelled LacI transcription factor when bound to *lacO* receptors on DNA in *E. coli*^42^ due to fluctuations of the DNA itself. We therefore denoted *D*_1_ as signifying static DNA-bound gyrase clusters. The “slow” diffusing component *D*_2_ with modal values 0.20 and 0.18 μm^2^/s for GyrA and GyrB respectively is consistent with earlier GyrA-only PALM tracking of clusters that are sliding on and/or transiently-bound to DNA, while the “fast” diffusing component *D*_3_ with modal values of 0.58 and 0.59 μm^2^/s for GyrA and GyrB respectively is consistent with clusters that freely diffuse in the cytosol. Although modal values of the three diffusion components were similar between GyrA and GyrB, the proportion of clusters associated with the static component is ∼20% higher for GyrB with ∼1/3 fewer clusters associated with the fast component (Fig. 2b insets).

### The mobility of gyrase clusters is lower in the presence of gyrase-targeting antibiotics

We applied two different classes of gyrase-targeting antibiotics, coumermycin A1 and ciprofloxacin, in Slimfield experiments at their minimum inhibitory concentrations (MIC) to investigate how gyrase clustering depends on its catalytic activity. Coumermycin A1 is an aminocoumarin that primarily targets the GyrB N-terminus by competitively inhibiting ATP binding^43^. The mean GyrA cluster diffusivity was reduced upon coumermycin A1 treatment to 0.29 ± 0.03 µm^2^/s (Fig. 2c, P= 4.6 × 10^−4^) the largest reduction for GyrA clusters colocalised to GyrB (Supplementary Fig. 6b). We measured no significant change in GyrB cluster diffusivity (P= 0.08). Applying the 3-component diffusion model showed that GyrA mobility reduction was mainly due to increases in both slow and static components (Fig. 2d). While the change in overall diffusivity of GyrB was not significant, there was a considerable increase in the static component (from 44% to 62%) at the expense of the slow component, indicating that coumermycin A1 had an effect on GyrB mobility as well.

The fluoroquinolone ciprofloxacin binds specifically to gyrase resulting in a stabilised DNA gyrase cleavage state; a nucleoprotein barrier to DNA replication that triggers cell death^44^ through generation of downstream double-strand breaks^45^. Using ciprofloxacin, GyrA cluster diffusivity was lowered (0.34 ± 0.03 µm^2^/s, Fig. 2e, P= 7.7 × 10^−3^), with the largest reduction for GyrA colocalised to GyrB (Supplementary Fig. 6c). We measure no significant difference to GyrB diffusivity (0.20 ± 0.02 µm^2^/s, Fig. 2e, P= 0.06). Applying the 3-component diffusion model showed that the reduction in GyrA mobility was mainly driven by an increase in the slow component (Fig. 2f). Similar to that observed for coumermycin A1, there was a considerable increase in the static component (from 44% to 68%) at the expense of the slow component on GyrB.

We compared these findings to those using cells treated with the antibiotic rifampicin^46,47^ which does not directly target gyrase but inhibits transcription via RNA polymerase. Our previous findings using a fluorescent GyrA reporter suggested that applying rifampicin resulted in a small increase in the fast component^26^ and only a moderate decrease in the slow/static components, so gyrase appeared to be performing its catalytic activity even when no (+) supercoils were added due to transcription. In our present study visualising GyrA and GyrB simultaneously, we measured no overall difference in GyrA diffusivity (0.54 ± 0.06 µm^2^/s, Fig. 2g, P = 0.63) compared to untreated cells. Similarly, the diffusivity of GyrB clusters (0.20 ± 0.04 µm^2^/s, P = 0.17) compared to untreated cells was not significantly different, irrespective of subunit colocalisation (Supplementary Fig. 6d). Applying the 3-component diffusion model indicated similar proportions of static, slow and fast to GyrA clusters from untreated cells, but showed that the observation of no overall change to mean GyrB mobility was due to a higher proportion of static offset by a lower proportion of slow GyrB diffusion; this suggests that transcription may still have an indirect effect on gyrase clusters mediated primarily through GyrB.

### ATP-binding inhibition increases GyrA and GyrB levels and clustering, whereas cleavage-complex trapping increases only GyrA

Measuring the cellular copy numbers and cluster stoichiometry values for cells treated with the two gyrase-targeting antibiotics indicated that GyrA copy number was significantly higher for coumermycin A1 treated cells (901 ± 39 GyrA per cell, P = 1.4 × 10^−17^), however, GyrB copy number showed no significant change (674 ± 32 GyrB per cell, Table 1 and Fig. 3a, P = 0.66). The mean stoichiometry for both GyrA and GyrB clusters was also significantly higher (Fig. 3b, Supplementary Fig. 4b) at 31.6 ± 1.1 GyrA (P= 4.6 × 10^−13^) and 61.2 ± 2.0 GyrB (P= 7.3 × 10^−5^). Coumermycin A1 was shown to lead to GyrB dimerisation by each molecule trapping two GyrB monomers^48^. This might explain the increased clustering of GyrB upon coumermycin A1 treatment.

**Figure 3.**
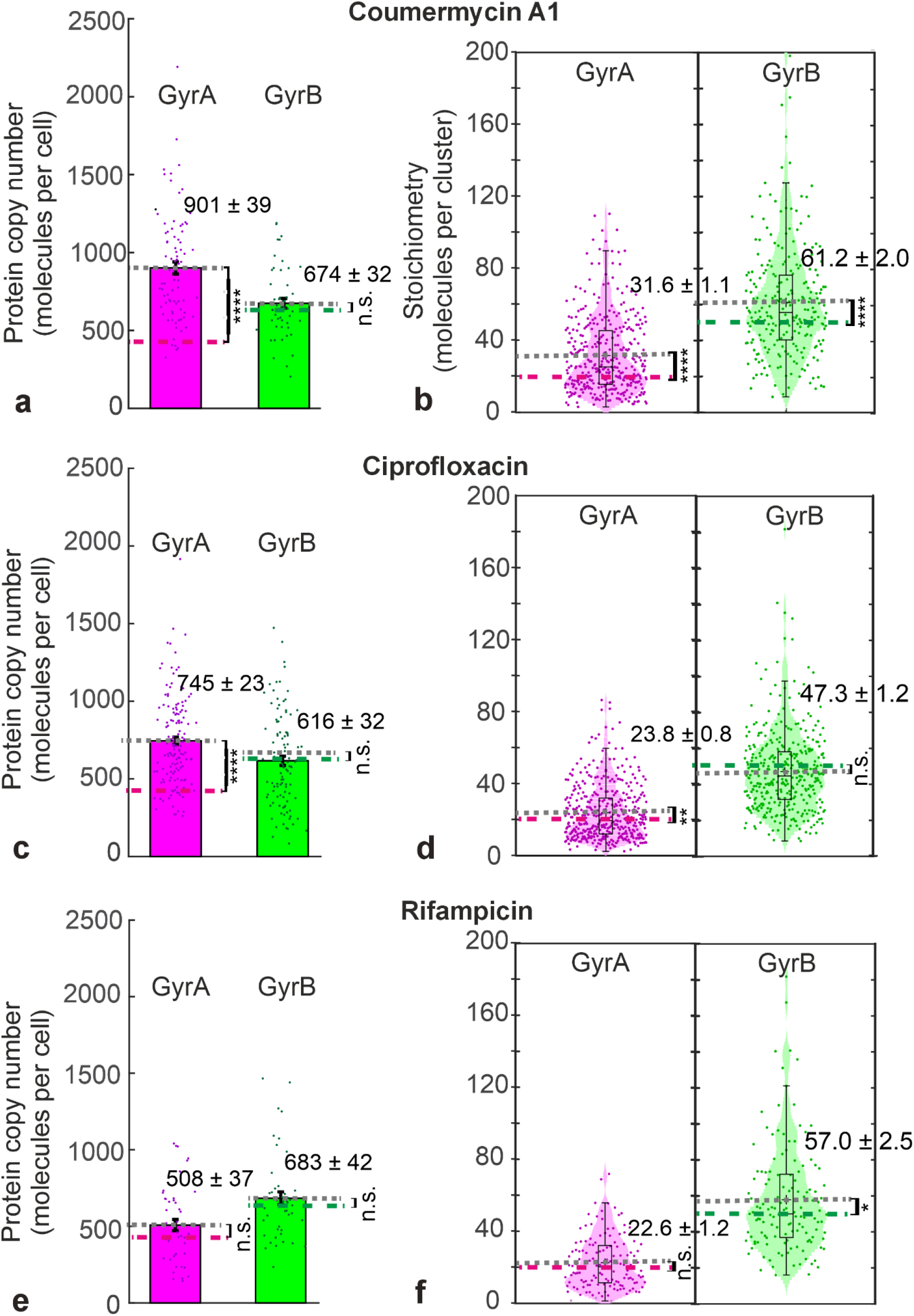
Bar charts with overlaid jitter data for copy number (left panels) and violin plots for cluster stoichiometry (right panels) for GyrA (magenta) and GyrB (green), stoichiometries, mean (grey dashed lines) and s.e.m (error bars) indicated in presence of **a.,b.**; coumermycin A1 (n = 366 GyrA clusters, n = 254 GyrB clusters); **d., e.** ciprofloxacin (n = 400 GyrA clusters, n = 375 GyrB clusters); **e.**, **f.** rifampicin (n = 140 GyrA clusters and 137 GyrB clusters); magenta and green dashed lines indicate mean values for untreated cells.

Treating cells with ciprofloxacin increased GyrA copy number to 745 ± 23 GyrA per cell (Table 1 and Fig. 3c, P = 3.8 × 10^−19^) and its mean stoichiometry (23.8 ± 0.8 GyrA, Fig. 3d, Supplementary Fig. 4c, P = 4.4 × 10^−3^). However, we measured no significant changes to either the copy number (P = 0.31) or stoichiometry in clusters (P = 0.12, Table 1) for GyrB.

Treating cells with rifampicin resulted in no significant changes to cellular copy numbers for GyrA (P = 0.40, not significant) or GyrB (P = 0.60, not significant, Table 1 and Fig. 3e). The mean stoichiometry values for GyrA and GyrB clusters in rifampicin-treated cells was not significantly different from those in untreated cells (Fig. 3f, Supplementary Fig. 4d, P = 0.11 and P =0.07).

Our observation of copy number sensitivity to gyrase-targeting antibiotics *in vivo* can be compared with earlier findings using multi-stage *in vitro* biochemical methods and antibody labelling of purified subunits to estimate protein expression levels of GyrA and GyrB in the presence of a range of antibiotics, including coumermycin A1 though not ciprofloxacin, suggesting upregulation of both subunits for relatively high antibiotic concentrations^12^. Our more direct measurements *in vivo* suggest that the sensitivity to gyrase-targeting antibiotics is more likely to involve GyrA, presumably because GyrB is already in excess. The physiological bottleneck upon inhibiting gyrase could therefore be the availability of sufficient GyrA to ensure catalytic activity, hence its upregulation.

### Gyrase clusters are liquid condensates

Since single-particle tracking and molecular counting for both GyrA and GyrB indicate that clusters are non-stoichiometric and do not exhibit a purely static mobility, we hypothesised that this could be due to liquid-liquid phase separation (LLPS). Over the past decade, multiple studies have identified LLPS as a driver to forming non-stoichiometric membraneless condensates in live cells including bacteria^49–52^, in a range of processes involving different protein and nucleic acid components. A hallmark of LLPS is the formation of weak, multivalent links that stabilise a liquid crystalline biomolecular assembly, which can be disrupted by application of the organic solvent 1-6 hexanediol (HEX)^51,53^ . Upon HEX treatment, we found that the proportion of GyrA colocalised to GyrB clusters (P = 0.21, not significant), and of GyrB colocalised to GyrA clusters (P = 0.23, not significant), drops only marginally between 6-9% (Fig. 4a). However, the cluster stoichiometry (Fig. 4b) for GyrA (13.5 ± 0.7 GyrA, P = 3.0 × 10^−6^, extremely significant) and GyrB (41.7 ± 1.7 GyrB, P = 3.0 × 10^−4^, highly significant) was reduced significantly. The reduction of stoichiometry, but not complete cluster disruption and only marginal reduction in GyrA/GyrB colocalisation, points towards HEX breaking weak liquid-like interactions involving a subpopulation GyrA and GyrB within clusters. Since gyrase heterotetramers are known to be bound to DNA via strong transient covalent interactions with GyrA, this observation points towards the HEX-sensitive population of GyrA and GyrB being those not strongly bound via covalent interaction to DNA.

**Figure 4.**
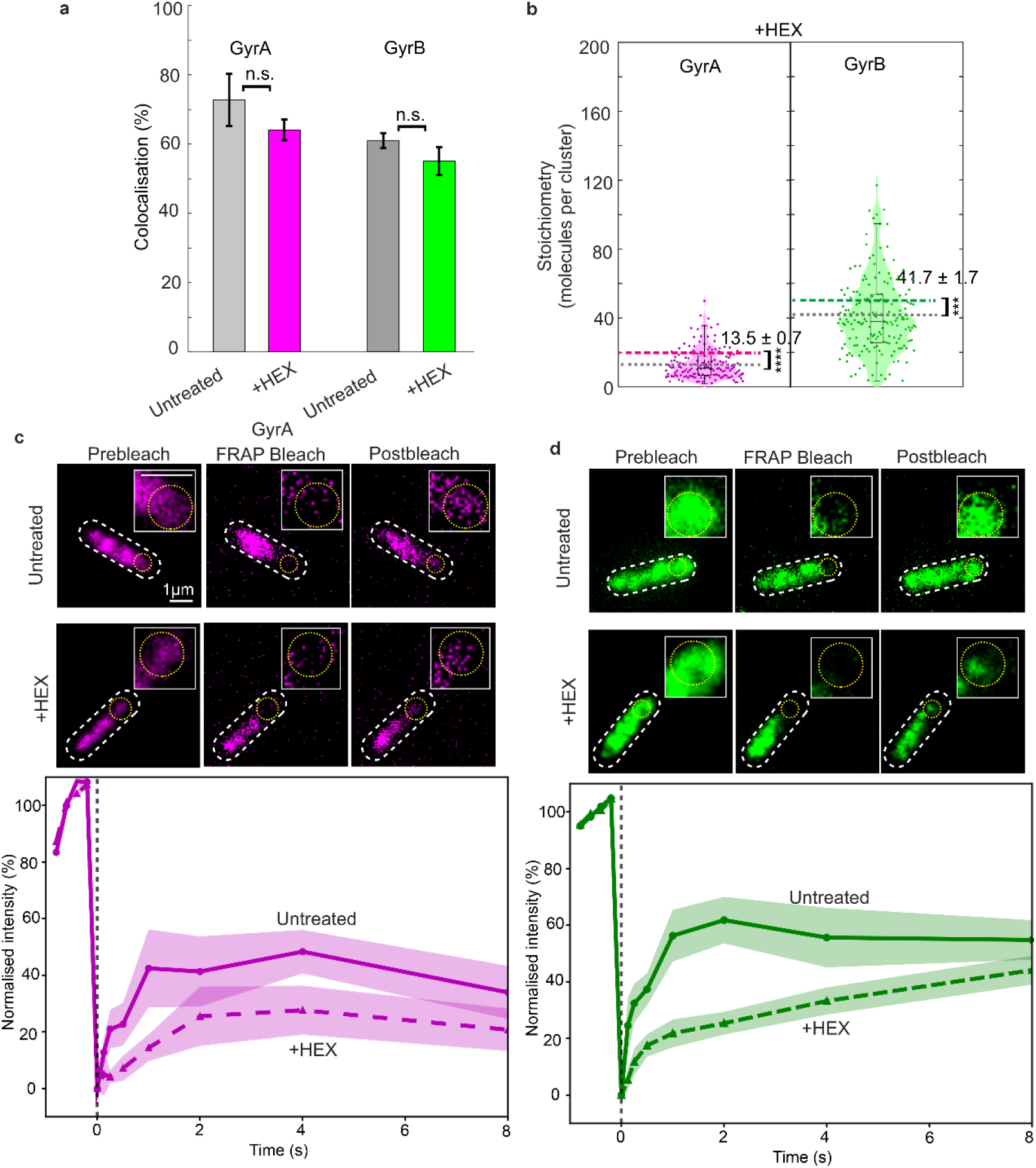
Chemical disruption of weak GyrA-GyrB interactions and FRAP indicate gyrase clusters have liquid properties. **a.** Colocalisation levels for GyrA (magenta) and GyrB (green) clusters treated with HEX compared against untreated cells (grey), s.e.m. error bars. Violin plots with jitter data overlaid for **b.** stoichiometry and **c.** diffusivity for HEX treated cells, mean levels (grey dashed lines) compared against equivalent untreated cells (magenta and green dashed lines. FRAP on **d.** GyrA and GyrB, upper panels are representative pre and postbleach images, extent of focused laser bleach indicated (yellow circle), lower panels showing normalised FRAP for untreated (solid lines) and HEX treated (dashed lines). Timepoint for focused laser bleach indicated by vertical black dashed line, s.e.m. shaded error bounds, number of FRAP traces in range 20-24.

Another approach used extensively to investigate kinetics of biomolecular LLPS condensates is fluorescence recovery after photobleaching (FRAP)^49^. We acquired four consecutive Slimfield images for each cell to establish the spatial pattern of GyrA and GyrB (Fig. 4c,d, top panel, prebleach images), then directed co-aligned 488 nm and 561 nm wavelength focused lasers to bleach a diffraction-limited region equivalent to ∼0.5 µm^2^ on 2D images (Fig. 4c,d, yellow circles) applied for ∼140 ms to completely photobleach mGFP or mCherry in that region. Fluorescence recovery was then monitored using dual-colour stroboscopic Slimfield over the subsequent 8 seconds. We observed loss of mCherry fluorescence in a cell region extending up to ∼0.5 µm beyond the initial focused laser bleach region, while that for mGFP was largely confined to the diffraction limited region (Fig. 4c,d, magenta and green zero-time postbleach panels). These observations are consistent with a higher mean mobility for GyrA compared to GyrB clusters (Fig. 2a) resulting in a greater likelihood of GyrA diffusing into the focused laser region during the bleaching period.

Fitting fluorescence data (Fig. 4c,d, lower panels, solid line traces) using a single exponential model indicated a lower steady-state level of recovery for GyrA of 42 ± 3% (± s.d.) compared to GyrB of 57 ± 3% (Supplementary Fig. 8 and Supplementary Table 3) with similar recovery half-times of 0.31 ± 0.11 s (± s.d.) and 0.24 ± 0.06 s for GyrA and GyrB respectively. Treating cells with HEX resulted in steady-state recovery levels of 25 ± 3% and 42 ± 5% for GyrA and GyrB respectively (Fig. 4c,d, lower panels, dashed line traces), equivalent to a reduction of 25-40% compared to untreated cells. HEX treatment resulted in recovery half-times of 0.70 ± 0.25 s and 1.38 ± 0.49 s for GyrA and GyrB respectively, a ∼2-5 fold relative increase (Supplementary Fig. 8 and Supplementary Table 3).

The decrease in steady-state recovery levels for GyrA and GyrB following HEX treatment although significant is only partial. This observation, when compounded with the reduction of GyrA and GyrB cluster stoichiometry following HEX treatment, points towards a sub-population of GyrA and GyrB clusters that interact weakly and non-covalently, which can be disrupted by HEX. GyrA and GyrB within clusters which are insensitive to HEX disruption are likely to be those bound to DNA via strong covalent interactions, consistent with extended *t_1/2_* turnover values. Taken as a whole, these observations indicate that gyrase clusters comprise a liquid-like component due to GyrA and GyrB that is not directly DNA bound via strong covalent bonds in addition to solid-like GyrA and GyrB directly and covalently bound to DNA. The discovery that the rate of turnover observed for GyrB following HEX treatment is lower than that of GyrA hints at a role for excess GyrB to maintain gyrase cluster integrity and, in effect, extend the processivity of gyrase on DNA.

### Excess GyrB in clusters forms multiple weak multivalent bridges between gyrase heterotetramers

To investigate the role of excess GyrB in maintaining gyrase cluster integrity and enhancing gyrase processivity on DNA, we performed structural docking analysis. Structures of individual gyrase GyrA and GyrB subunits from PDB ID: 9GBV^54^ (domain residues indicated in Fig. 5a) were clustered and evaluated using an empirical energetic docking score, with a more negative value indicating a more likely attractive interaction^55–59^. The confidence score was calculated from the docking score, with a value of 0.7 or higher indicating very high probability of binding. We docked individual gyrase subunits with themselves, observing that GyrA and GyrB bound together with the highest confidence score (0.9998) compared to any other combination of single subunits (Fig. 5b and Supplementary Table 4). Two docked GyrBs had a lower confidence score (0.859) compared to two GyrAs (0.998). Two of the A:B dimers were then docked to form the heterotetramer with a confidence score of 0.996, which is higher than that obtained by docking the two homodimers B:B meaning a higher likelihood of the first docking scenario occurring *in vivo*. As these confidence scores were very high, it was assumed that these would occur early in the clustering process and therefore the heterotetramer (2(GyrA+GyrB)) was used as a sensible starting point to build up a gyrase cluster with the corresponding relative stoichiometry comparable to those found in clusters *in vivo*. It was generated by docking two further GyrBs, followed by another heterotetramer and two more GyrBs. The summary of the docking interactions is illustrated in Fig. 5c and Supplementary Movies 1-5 which culminates in a final gyrase cluster comprising two A_2_:B_2_ heterotetramers and four excess GyrB monomers; the docking algorithm used can be extended to generate arbitrary sized clusters while still maintaining the same approximate relative stoichiometry between GyrA and GyrB. The fact that the confidence scores for all the docking steps are higher than 0.7 indicates that these interactions are highly likely to occur.

**Figure 5.**
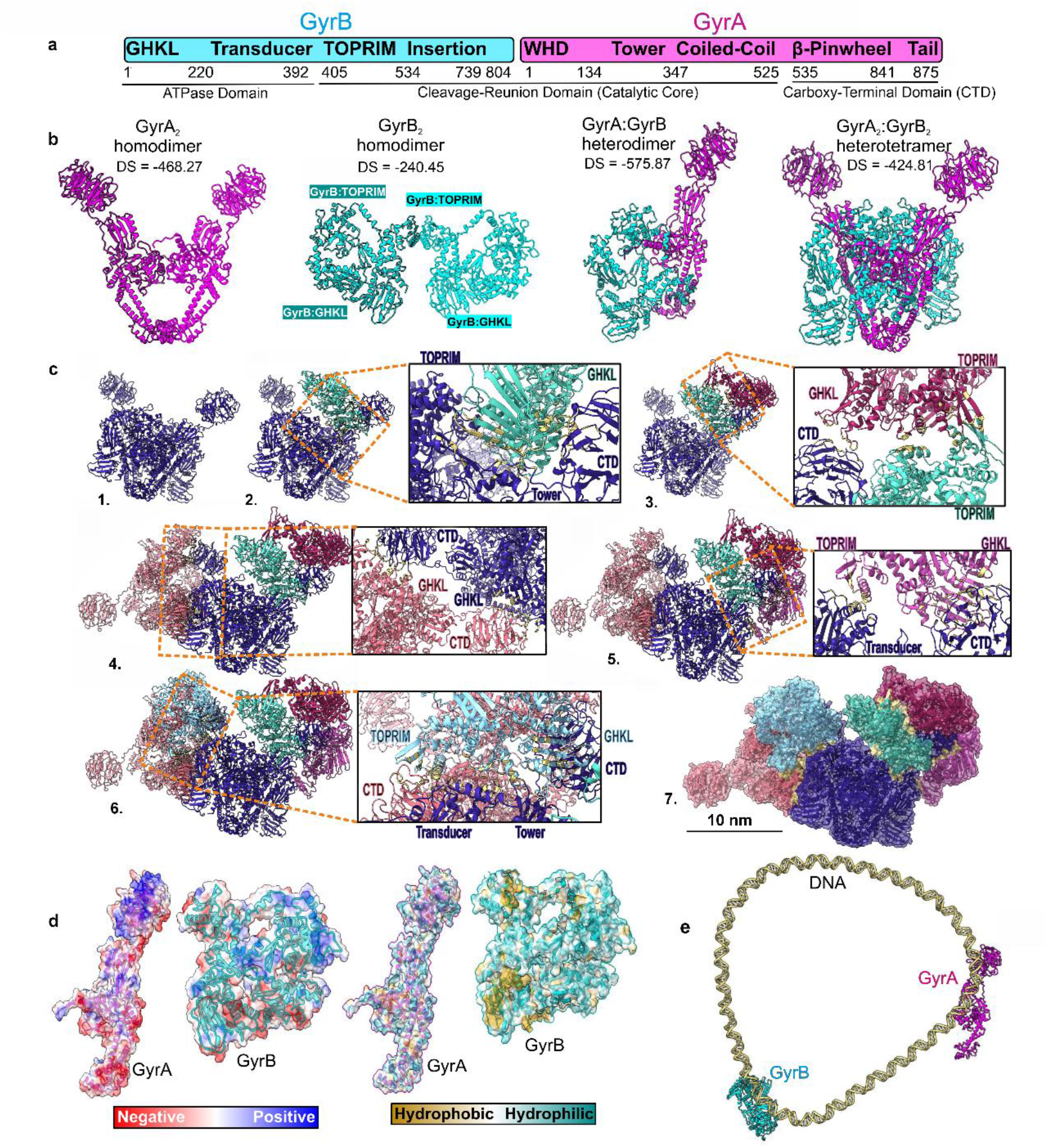
Structural docking suggests that excess GyrB interacts with gyrase heterotetramers. **a.** Residue numbers and domains of each gyrase subunit from PDB ID 9GBV^60^. **b.** Docking structures using GyrA (magenta) and GyrB (cyan) with docking scores (DS) indicated. **c.** Stepwise building of gyrase cluster with relative stoichiometry A:B=1:2, zoom-in panels (orange dash) showing interaction interfaces, with buried surface area residues shown (yellow): **1**. Heterotetramer (dark blue) used as seed structure for building cluster; **2.** GyrB added (green); **3.** Second GyrB added (dark pink) interacting with heterotetramer and first GyrB; **4**. Second heterotetramer added (light pink) interacting with first heterotetramer; **5.** Third GyrB added (purple) interacting with first heterotetramer; **6.** Fourth GyrB added (light blue) interacting with both heterotetramers; **7.** Final cluster with surfaces and interaction interfaces indicated. **d.** Electrostatic and hydrophobic surfaces on individual gyrase subunits. **e.** Docking gyrase in the presence of DNA, average structure from a simulation on a relaxed DNA minicircle of 339 bp^61^ was used as a substrate (offering a range of curvatures that could be selected to optimize the docking score) showing GyrA (magenta) and GyrB (cyan) separately docked to DNA.

We found that the added excess GyrB could interact with tetramers through additional surface areas being able to establish weak multivalent interactions between them. The different types of the residues involved in these interfaces indicates the diverse nature of contacts stabilizing these clusters, which include hydrogen bonding as well as electrostatic, hydrophobic and hydrophilic interactions (Fig. 5d, Supplementary Fig. 9 and Supplementary Movies 1-5). Nevertheless, a surface map of the electrostatic potential on both subunits show that GyrB has a more neutrally charged surface than GyrA, which includes clusters of positive and negative residues. In terms of hydrophobicity, GyrB is predominantly hydrophilic with a few hydrophobic pockets, whereas GyrA is more neutral. The findings show that the primary nature of the interactions governing these clusters would vary on the encountered subunits, being largely electrostatic between two GyrA, hydrophilic/hydrophobic between two GyrBs, and polar-electrostatic between a GyrA and a GyrB. Attempting docking with both GyrA and GyrB added subunits resulted in a less negative docking score compared to docking just with GyrB (Supplementary Table 4) resulting in filamentous as opposed to clustered structures for which we saw no direct evidence experimentally (Supplementary Fig. 10). We find therefore that the docking analysis is consistent with GyrB being the primary bridging component between heterotetramers.

To elucidate why GyrA exhibits greater mobility than GyrB within cells, docking simulations were conducted for gyrase subunits on DNA (Supplementary Table 5, example of docking of monomeric GyrA and GyrB to DNA shown in Fig. 5e). The average structure from a simulation on a relaxed DNA minicircle of 339 bp was chosen as a substrate, as it offered a variety of curvatures that could be selected to optimize the docking score. The GyrB monomer or homodimer formed a cluster with DNA exhibiting a greater confidence score than GyrA monomers or homodimers, corroborating experimental findings that suggest GyrB has lower diffusion rates than GyrA. When examining the interactions responsible for this binding, we found that the presence of polar residues in GyrB but the absence of negatively charged residues in GyrA allow the protein to establish favourable contacts with the highly negatively charged DNA molecule.

## Discussion

A principal finding from this work is that DNA gyrase can occur in complexes that contain super-stoichiometric amounts of GyrA and GyrB and that such complexes may have a functional significance in terms of accelerated ability to carry out DNA supercoiling. Following discovery of gyrase in *E. coli* by Gellert and co-workers^62^, it was found to consist of two subunits, now called GyrA and GyrB^63–66^. It was later established, using gyrase purified from *E. coli* and *Micrococcus luteus*, that the active enzyme likely consists of an A_2_B_2_ complex^67,68^ although in other species, A_2_ and A_2_B, were also detected^68^. In the ensuing years, evidence for other complexes has been found. GyrA is generally a dimer in solution^68,69^, although monomers can be detected by mass spectrometry^70^. Higher-order oligomers of GyrA have also been reported, e.g., in *Mycobacterium tuberculosis* and *M. smegmatis*, GyrA tetramers (A_4_) were found^71^. In *Bacillus subtilis* the GyrA NTD (N-terminal domain) can form hexamers^72^. Using mass spectrometry monomers, dimers (A_2_), tetramers (A_4_) and hexamers (A_6_) of *E. coli* GyrA have been detected^70^. Using cryo-EM, both dihedral and tetrahedral complexes of *Streptococcus pneumoniae* GyrA-NTD were observed in the presence of DNA^73^. The dihedral complex, built from 4 GyrA-NTD dimers, predominates when GyrB is excluded; addition of GyrB and novobiocin promotes formation of the tetrahedral complex, built from 6 GyrA-NTD dimers, but does not contain GyrB^73^.

In contrast, it has been suggested that GyrB is largely a monomer with the capacity to form dimers^74^; dimerization is favoured by binding of ATP or non-hydrolysable nucleotide ADPNP (5’-adenylyl-β,γ-imidodiphosphate)^21,75^. However, recently, multimers of *E. coli* GyrB (tetramers and beyond) have been found using native gel electrophoresis^70^; furthermore, GyrB octamers and higher-order aggregates have been observed using mass spectrometry^76^.

Multimerisation has been reported in other type II topoisomerases, for example, yeast topo II has been shown to form multimers (tetramers and beyond) that may have a role in chromatin structure^77^; however it remains to be seen whether higher-order complexes of gyrase are of functional significance. Our present work suggests that higher-order gyrase molecular assemblies might function in resolving (+) DNA supercoils. A key role of gyrase is that it removes (+) DNA supercoils ahead of advancing replication forks during bacterial DNA replication^3^. One conundrum is that bacterial replication (e.g. in *E. coli*) can proceed at ∼1,000 nucleotides per sec, whereas if the rate of (−) supercoiling by gyrase is limited by its rate of ATP hydrolysis (∼1 per sec)^23,65^ this would require ∼50 gyrase molecules per advancing fork. One previous study suggested^20^ that gyrase might relax (+) supercoils at burst rates of ∼100 per sec; however, it is not clear in this work whether this entails very rapid rates of ATP hydrolysis, somehow overcoming the normal rate-limiting step, the release of ADP^21^. One possibility that has emerged from our present study is that macromolecular gyrase assemblies comprising super-stoichiometric amounts of GyrB could circumvent this problem.

Another proposed function of gyrase higher-order assemblies is in illegitimate recombination (IR). This refers to recombination between non-homologous sequences of short regions of homology on two different DNA molecules or at two different sites of a DNA molecule and results in deletions, tandem duplications insertions or inversions^78,79^. IR is proposed to be mediated by gyrase and has been shown to be stimulated by quinolones such as oxolinic acid and can be abolished by the aminocoumarin drug coumermycin A1^80^. Although the exact mechanism is not known, it has been proposed to involve two gyrases on adjacent pieces of DNA forming a transient complex and subunit exchange and concomitant exchange of DNA ends^81^.

Gyrase higher-order assemblies could also provide an explanation for the observed ability to catalyse supercoiling when it contains a single catalytic tyrosine; it has been suggested that this might occur via a swivel (nicking-closing) mechanism distinct from the widely accepted double-strand passage mechanism common to type II topoisomerases^82^. An alternative explanation is the possibility that the enzyme undergoes subunit exchange (as above for IR) or interface swapping, involving reconstitution of a DNA gate with two tyrosines^70^, with our observations of turnover of subunits within clusters offering support to this hypothesis.

Previous *in vitro* titration measurements for gyrase subunits in *B. subtilis* and *E. coli*^83^ suggested that GyrA only reaches stoichiometric saturation with GyrB when the latter is added in excess of a 1:1 A:B ratio (this work reporting 1:4) though with no direct experimental indication as to the cause. Our present work in live cells offers insights to this observation, though our biochemical findings from EMSA *in vitro* indicate that an excess of GyrB in a mixture of GyrA and GyrB does not result in significant differences to the level of binding to a 167 bp DNA fragment. Using high precision bioimaging of GyrA and GyrB we observe non-stoichiometric gyrase molecular assemblies which demonstrate a range of relative A:B stoichiometries as opposed to a single fixed value, but all indicating an excess of GyrB, with a mean ratio of 1:2.5. We find that these variable relative stoichiometries can be significantly biased by treating cells with gyrase-specific antibiotics, indicating a sensitive coupling between cluster mobility and stoichiometry and the catalytic activity of the gyrase enzyme.

Curiously, treatment with both ciprofloxacin and coumermycin A1 resulted in upregulation of only GyrA and not GyrB. The stoichiometry of GyrA clusters was also found to increase upon treatment with both antibiotics. Only coumermycin A1 treatment increased clustering of GyrB which might be explained from an earlier observation that each molecule of the antibiotic binds two monomers of GyrB^48^. However, this was not accompanied with any increase in GyrB copy number. GyrA is known to be trapped mid-catalysis upon ciprofloxacin treatment which might explain the increased expression and clustering of GyrA. The gyrase cycle is also clearly stalled by coumermycin A1, however, since extensive biochemistry and structural biology data indicate that its mode of inhibition is very different to that of ciprofloxacin, the cause for this catalytic stalling remains an intriguing open question.

Our findings that GyrA and GyrB turnover within clusters, coupled to observations that gyrase clusters are disrupted by treatment with HEX, support a biomolecular condensate model. Condensates have previously been implicated in topoisomerase function in modulating the catalytic activity of eukaryotic topoisomerase II^84^. Here, we find that gyrase condensate stability is mediated through multiple weak interactions involving excess GyrB (Fig. 6). Structural docking shows that these interactions involve multivalent bridging between GyrB and A_2_B_2_ heterotetramers and chimes with earlier observations *in vitro* that functional reconstituted gyrase requires excess GyrB^83^ to relax positively supercoiled DNA^83^. One outcome of such metastability is that subunit turnover ensures that gyrase clusters can form and disassemble dynamically depending on local requirements set by the state of (+) supercoiling on the genome, whilst still enhancing the effective processivity of the gyrase enzyme on the DNA through the adhesive forces of a liquid condensate: if an individual gyrase heterotetramer spontaneously dissociates from DNA then, if it is not confined within a condensate, it may be free to diffuse away; however, within a condensate, such diffusion is counteracted by surface tension which thus increases the chances that unbound heterotetramers rebind DNA and thus can engage in multiple subsequent catalytic cycles. This condensate model potentially reconciles a longstanding conundrum for how gyrase in live bacteria keeps up with rapid accumulation of (+) supercoils due to DNA replication. Future work, beyond the scope of our current study, will be needed to quantify if the *in vivo* clustering of gyrase increases its effective catalytic rate by the ∼3 orders of magnitude predicted.

**Figure 6.**
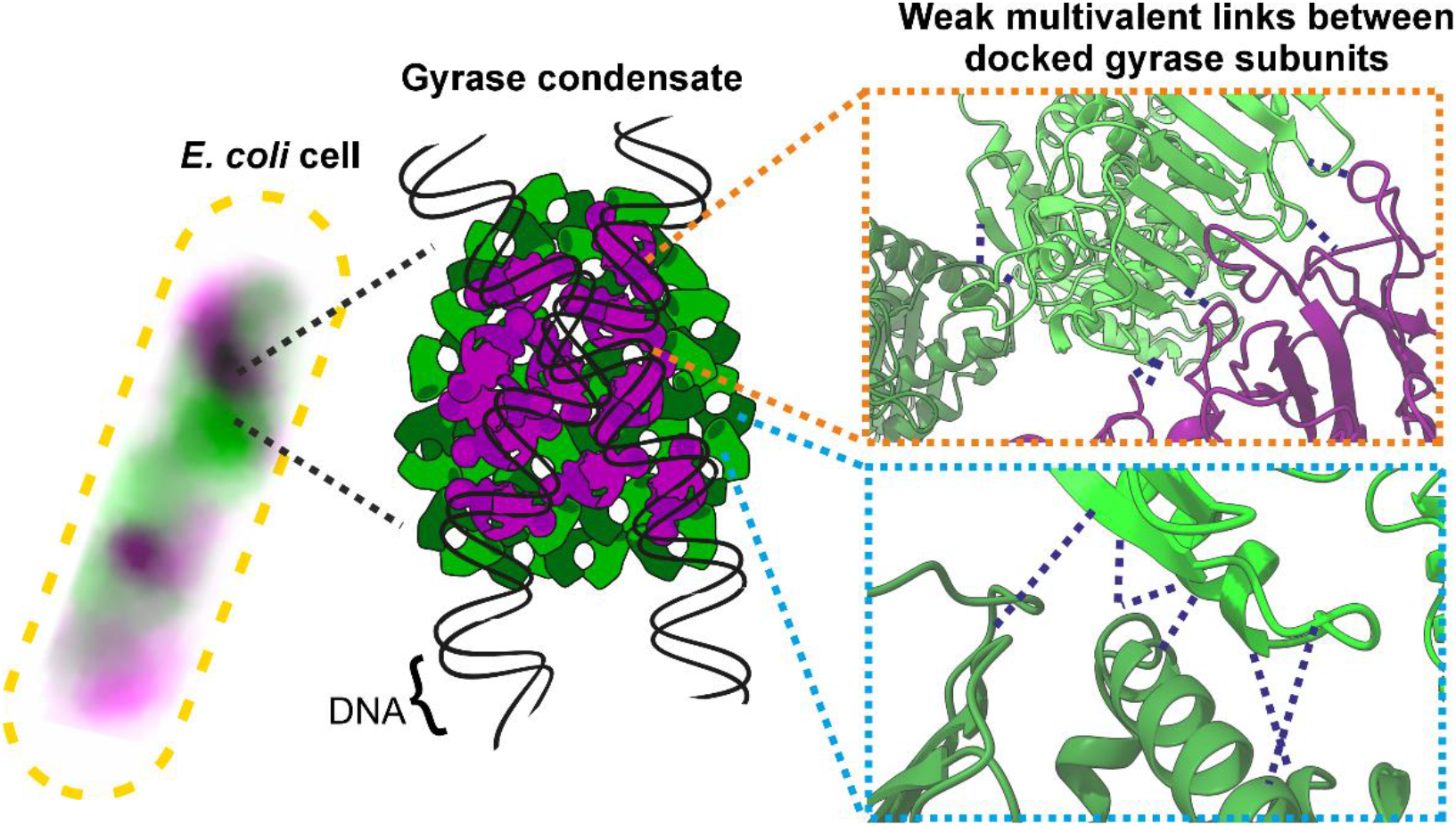
Excess GyrB stabilises DNA gyrase liquid condensates bound to DNA. Cartoon model illustrating the presence of liquid gyrase condensates bound to DNA which are stabilised through weak multivalent interactions between GyrA (magenta) or GyrB (green) incorporated in A_2_B_2_ heteroteramers, and excess monomeric GyrB. Dashed dark blue lines indicate predictions from ChimeraX for hydrogen bonding between docked gyrase subunits.

Our findings shed light on the fundamental question of how bacteria resolve potentially lethal accumulation of DNA (+) supercoils. We have discovered that non-stoichiometric molecular assemblies formed from heterotetramers and excess GyrB can offer a mechanism to leverage the biophysical metastability of liquid condensates to facilitate enhancement of catalytic activity above the level of isolated gyrase heterotetramers bound to DNA. Gyrase is essential to bacteria and is thus an attractive target for novel antibiotic development; our new insights of the condensate properties of gyrase *in vivo* is an important step forward in understanding the mechanistic actions of this enzyme in bacterial metabolism and in designing novel therapeutics to potentially target gyrase condensate disruption. These findings demonstrate that, beyond their primary association with spatial compartmentalisation, signalling and regulation, biomolecular condensates might directly enhance the catalytic efficiency of an essential molecular machine by increasing local enzyme recycling. This concept may have implications for understanding how dynamic molecular assemblies throughout biology overcome kinetic limitations imposed by diffusion.

## Methods

### Strain construction and characterisation

Strains containing *gyrA-mCherry* and *mGFP-gyrB* used are derivatives of laboratory wild type strain TB28. Briefly, for tagging *gyrA*, *linker-mCherry* followed by a kanamycin resistance cassette flanked by *frt* sites was amplified by PCR from the plasmid pJGB374^29^ using primers oAS182 and oAS183 (Supplementary Tables 6 and 8). Amplification primers had a 50 bp homology at their 5’ end to the last 50 bp of the *gyrA* gene preceding the stop codon (forward primer) or the 50 bp immediately after the stop codon (reverse primer). Resulting PCR products could thus recombine with the chromosome to produce in-frame integration of *linker-mCherry-<kan>* immediately downstream of *gyrA*, resulting in the *gyrA-mCherry-<kan>* allele.

PCR products were first treated with *Dpn*I and then gel purified. The purified PCR products were introduced into cells expressing the lambda Red genes from the plasmid pKD46^85^ by electroporation. The recombinants were selected for kanamycin resistance and screened for ampicillin sensitivity. The colonies obtained were verified for integration by PCR with primers oJL1 and oJL14. The PCR products were then confirmed by sequencing using primers oJGB398, oJGB399, oMKG71, oMKG72, oJL1 and oJL14.

To generate the *<kan>-mGFP-gyrB* allele, the *<kan>-mGFP-linker* cassette was amplified from plasmid pAP1 using primers oAS195 and oAS196. The oAS195 primer had 50 bp homology immediately upstream of *gyrB* while oAS196 had 50 bp homology to the first 50 bp of the *gyrB* ORF excluding the start codon. The amplified product would thus recombine on the chromosome such that the fusion would generate the *<kan>-mGFP-gyrB* allele under the control of the native *gyrB* promoter. The PCR product was introduced on the chromosome of cells expressing lambda red genes after *Dpn*I digestion, gel extraction, and electroporation as described above for the *gyrA* fusion. The recombinants were verified by PCR amplification using primers oJL15 and oAS188 and confirmed by sequencing the PCR products with oJL15, oMKG70, oMKG71, oMKG72, oJGB417 and oJGB418.

The kanamycin-resistance gene from *<kan>-mGFP-gyrB* was removed by expressing Flp recombinase from plasmid pCP20^85^ to generate the kanamycin-sensitive strain carrying the FP fusion allele *<>-mGFP-gyrB*. The *gyrA-mCherry-<kan> <>-mGFP-gyrB* dual labelled strain was created by introducing the *gyrA-mCherry-<kan>* allele by standard P1-mediated transduction into the single labelled *<>-mGFP-gyrB* strain.

### Growth curves

Strains were grown overnight in LB medium at 37°C at 200 rpm. Overnight cultures in stationary phase were diluted the next morning 100-fold into fresh LB or washed once with M9 medium and diluted 100-fold in fresh M9 medium supplemented with 0.2% glycerol as the carbon source. Aliquots of 100 μL from the diluted cultures were dispensed into individual wells of 96-well clear flat-bottom sterile microplates (Corning). The microplate containing the diluted cultures was incubated in a BMG LABTECH SPECTROstar Nano microplate reader at 37°C and the optical density (A_600_) was recorded at defined time intervals. The time taken for the optical density to double during the exponential growth phase was taken as the generation time. The values expressed are the means of three independent replicates, with the standard deviation and errors indicated.

Based on previous characterisation, strains had a 10-15% content^32,86^ (i.e. immature non-photoactive fluorescent protein). All plasmids used in strain construction are indicated in Supplementary Table 6, strains used in this study are indicated in Supplementary Table 7 and primers in Supplementary Table 8.

### Testing minimal inhibitory concentration (MIC) of antibiotics

Antibiotic sensitivity was assessed using spots tests. The dual fluorescent protein labelled strain was grown overnight in LB at 37°C at 180 RPM to stationary phase. Serial 10-fold dilutions were then spotted onto LB plates with various antibiotics at the indicated concentrations. The isogenic wild type parent TB28 was included as a control and spotted similarly. The two strains were also spotted on LB only plates (0 μg/ml antibiotic) as a permissible growth control. MIC values for the fluorescently labelled strain were measured for the fluoroquinolone ciprofloxacin (MIC 10 ng/ml), the aminocoumarin coumermycin A1 (MIC 10 μg/ml) and the transcriptional inhibitor rifampicin, which does not directly target DNA gyrase (MIC 50 μg/ml) (Supplementary Fig. 2).

### Protein purification and supercoiling assays

Fluorescently labelled GyrA and GyrB subunits were expressed and purified separately. Both were expressed with poly-histidine N-terminal tags that could be cleaved. *E. coli* Rosetta2 cells were transformed with the relevant plasmid, grown to A_600_ of ∼0.6 and induced at 30°C for 3 hours. Cells were harvested, resuspended in 50 mM Tris·HCl (pH 7.5), glycerol (10% v/v), 1 M NaCl (buffer TGN) and lysed by passage through an Avestin cell disruptor. After centrifugation, the supernatant was loaded onto a 5 ml Ni-IMAC column in the same buffer and eluted with a gradient of imidazole in TGN. Fractions were analysed by SDS-PAGE and those containing the protein of interest were dialysed versus 50 mM Tris·HCl (pH 7.5), glycerol (10% v/v), 1 mM EDTA (buffer TGE) and then the histidine tag was removed by addition of TEV protease. This leaves a single additional glycine residue on the N-terminal of the proteins. After overnight digestion the protein was dialysed versus TGN buffer and loaded back onto the 5 mL Ni-IMAC column and the flow-through containing the cleaved protein was collected. This was dialysed into buffer TGE containing 1 mM DTT (TGED) and loaded onto a HiResQ column equilibrated in the same buffer. It was then eluted with a gradient to 1 M NaCl in TGED. Fractions containing the protein were pooled, dialysed versus storage buffer and stored at -80°C.

The subunits were mixed in a 1:1 ratio as far as possible and this was used in subsequent supercoiling assays. Essentially these consisted of a mix of 6 µl of 5X assay buffer (final 1X composition equivalent to 35 mM Tris·HCl (pH 7.5), 24 mM KCl, 4 mM MgCl_2_, 2 mM DTT, 1.8 mM spermidine, 1 mM ATP, 6.5% (w/v) glycerol, and 0.1 mg/mL albumin), 0.5 µl of relaxed pBR322 (1 mg/mL) and water plus either dilution buffer (50 mM Tris·HCl (pH 7.5), 100 mM KCl, 2 mM DTT, 1 mM EDTA, and 50% (w/v) glycerol), and the individual subunits to check for background activity or gyrase dilutions in a total volume of 30 µL. After incubation at 37°C for 30 min the reactions were stopped by the addition of 15 µL of STEB (40% (w/v) glycerol, 100 mM Tris·HCl (pH 8.0), 100 mM EDTA, 0.5 mg/mL Bromophenol Blue) and 30 µl of chloroform/isoamylalcohol. They were briefly vortexed ∼5 secs and centrifuged for 1 minute, and 20 µL of the upper aqueous layer loaded onto a 1% (w/v) agarose TAE gel. After electrophoresis (90 V for 90 minutes in TAE: 40 mM Tris·HCl (pH 8.3), 20 mM acetate, 1 mM EDTA) the gel was stained with 1 μg/mL ethidium bromide in water (15 mins), destained (5-10 mins) in TAE and visualised with a gel transilluminator. The appearance of a negatively supercoiled band on the gel at increasing intensity with increasing mixture volumes was clearly indicative of functional supercoiling activity.

### Gyrase-DNA electrophoretic mobility shift assays (EMSA)

GyrA:GyrB tetramers were reconstituted by combining purified GyrA purified as described previously^54^ with GyrB (Inspiralis) to generate either a 1:1 tetramer (1 μM GyrA and 1 μM GyrB) or a 1:5 tetramer (1 μM GyrA and 5 μM GyrB), each at a final tetramer concentration of 500 nM. Tetramers were prepared in tetramer dilution buffer containing 100 mM potassium glutamate, 50 mM Tris-HCl (pH 7.5), 2 mM DTT, 5 mM MgCl_2_, and 10% (v/v) glycerol. Electrophoretic Mobility Shift Assays (EMSA) were performed using a Cy5-labelled 167 bp DNA fragment (1 pmol in each lane). For each sample (1-6), GyrA and GyrB were pre-mixed at either 1:1 or 1:5 molar ratio, followed by incubation with DNA for 1 h at 37 C in 0.1 mg/ml BSA, 40 mM potassium glutamate, 25 mM Tris-HCl (pH 7.5), 2 mM DTT, 5 mM MgCl₂, and 9% (v/v) glycerol, 20 µl total volume. The reaction mixture was loaded onto a 5% agarose gel, electrophoresed at 110 V for 45 min, and imaged for Cy5 fluorescence. At matched GyrA amounts, increasing GyrB 5-fold (lanes 2 vs 4, or 3 vs 5 in Supplementary Fig. 5) did not result in measurable change of the proportion of gyrase-bound DNA to within ∼5% based on quantification of the scanned band intensities.

### Preparation of cells for imaging

Strains were streaked onto LB plates, containing antibiotics as appropriate, and incubated at 37°C overnight. Single colonies were inoculated into LB media and grown at 37°C shaking at 180 rpm for 5 hours. The culture was used to inoculate M9 minimal medium (1:50) supplemented with 0.2% glycerol and grown overnight at 37°C to *A*_600_ of 0.4–0.6. Next morning, the cultures were diluted 10 times into fresh M9 glycerol and grown again to A_600_ of 0.4–0.6. The cells were centrifuged and immobilized for imaging on 1% agarose pads containing M9-glycerol growth medium. Where indicated cells were incubated with either the appropriate antibiotic at its relevant MIC for 30 min, or with 5% HEX, for 5 min in liquid culture prior to imaging, with the equivalent concentration of antibiotic or HEX present during imaging.

### Slimfield microscopy

To visualise the fluorescent reporters of GyrA and GyrB in live *E. coli* we used a custom-built dual-colour single-molecule Slimfield microscope. Brightfield and fluorescent images were acquired using a custom OpenFrame (Cairn Research) with samples mounted on a precision nanostage (Mad City Labs) and a Prime95B sCMOS camera (Teledyne Photometrics) in 12-bit ‘Sensitivity’ or 16-bit ‘High Dynamic Range’ linear gain modes. A 100X oil immersion objective (NA 1.49 Apo TIRF, Nikon) was used with an additional 2.2X lens relay for a total magnification of 220X at the detector, resulting in a pixel size in Slimfield image sequences of 53 ± 5 nm. Prior to the detector, green and red channels were split using a bespoke colour splitter consisting of GFP/mScarlet-I dichroic beamsplitter (Chroma ZT488/561rpc) and bandpass emission filters (525/50 nm and 594/25 nm respectively). Laser excitation was produced from 488 nm and 561 nm continuous wave lasers (Coherent Obis LS) coupled to the microscope body at 20 mW and 40 mW source power respectively, then reduced in the sample plane to an FWHM of 25 µm, and vignetted to 10 µm diameter, resulting in mean excitation intensity of ∼1 kW/cm^2^. Alternating laser excitation between odd and even frames at 2 ms time intervals was produced by digitally modulating the output via the camera exposure trigger.

### Single particle tracking and diffusion analysis

Slimfield data were analysed using bespoke MATLAB and Python software as described previously^30,87^, with the signal to noise ratio set to >0.2. The intensities of single mGFP or mCherry molecules was determined from stepwise photobleaching analysis of overtracked foci beyond their bleaching point. These intensities were used to determine the stoichiometry of the corresponding fluorescent foci by dividing the observed intensity with that of the single fluorophore. Colocalisation between foci was determined as described earlier^36,88^. The microscopic diffusion coefficients were determined as described previously. Briefly, diffusion coefficients (*D*) for the detected foci tracks were determined by fitting a straight line to the initial component corresponding to the first three consecutive mean square displacement (MSD) datapoints of each track in addition to a fourth virtual datapoint given by 2σ^2^ where σ is the 1-dimensional measured localization precision of 40 nm.

Gamma fits followed the protocol of reference^26^ but instead of independently fitting the diffusivity histogram for each treatment to extract the diffusion coefficients for each population, tracks from across all treatments were collated and fit to a sum of one up to four gamma probability density functions, with a shape factor of four. The bin width of the histogram and the position of the ‘static’ peak were fixed at 0.045 μm/s^2^ which represents the experimental error in the calculated diffusion coefficient. Statistical analysis using the Fisher F-test indicated that to account for the combined distribution of diffusion coefficients for both GyrA and GyrB clusters minimally required the weighted sum of three distinct gamma probability density functions comprising a static, slow and fast diffusive population with diffusion coefficients *D_1_*, *D_2_* and *D_3_* respectively. The proportion of tracks in each treatment belonging to each population was determined by integrating the area under the diffusivity histogram in the three regions defined by the crossing points of three gamma fits: x1 between *D_1_*, *D_2_*, and x2 between *D_2_*, *D_3_*. Then proportion one is n<x1, proportion 2 is <x2 but >x1, proportion 3 is >x3.

### FRAP experiments

FRAP data were collected using the same bespoke Slimfield microscope as before but using the quarter waveplate to circularly polarize incident light before splitting the horizontal and vertical components into a narrowfield^89^ epifluorescence path and a focused confocal path via a polarizing beamsplitter such that 50% of the power is distributed to each path. The beam was then expanded such that the back aperture of the objective lens was just overfilled, generating a diffraction-limited excitation volume at the focal plane. FRAP acquisitions were captured by first recording four frames of Slimfield at 20 mW and 40 mW source power for mGFP or mCherry imaging respectively, and 4.6 ms exposure time. Then the shutter to the FRAP path was opened to photobleach one end of the cell for 30 consecutive frames, totalling 138 ms. Subsequent Slimfield frames were recorded at 0.125 s, 0.25 s, 0.5 s, 1s, 2 s, 4 s and 8 s to record the time dependence of fluorescence recovery.

For analysis, a region of interest (ROI) was drawn to define the area of the cell using ImageJ, a separate ROI was then drawn to define the area of the cell which was photobleached by the confocal laser. The fluorescence recovery was calculated as the mean intensity of the pixels inside the bleached region divided by the mean intensity of the cell after subtraction of the background signal to account for photobleaching. The background was calculated as the total intensity in the bleached ROI in the first post-bleach frame. Traces were normalized to the mean intensity of the cell during the initial four pre-bleach frames of Slimfield. Traces for each condition were averaged onto a mean recovery curve, which was fitted to an exponential solution for the reaction-diffusion equation^31^ using Scipy curve fit:

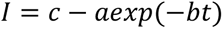

Where *a* is the amplitude of recovery, *b* is the rate of unbinding (1/*k_off_*), and *c* is the asymptote for recovery. The Shapiro-Wilks test was used as a goodness of fit metric by testing the normality of the residuals.

### Structural docking

The HDOCK server^55–59^ was used to cluster PDBs of individual GyrA and GyrB subunits^54^. One GyrA and one GyrB subunit were docked with high affinity -with a confidence score of 0.9998, which was higher than any other combination of single subunits. Two of these dimers were then docked to form the heterotetramer with a confidence score of 0.996, meaning a much higher likelihood of the docking scenario occurring *in vivo*. As these scores were so high, it was assumed that these would occur before much of the clustering and so the heterotetramer (2(A+B)) was used as a starting point to build up a cluster with the corresponding stoichiometry to those found in clusters *in vivo* (A:B ≈ 1:2). It was built up by adding two GyrBs to this, and then another heterotetramer and two more GyrBs. Each step was visualised with UCSF ChimeraX^90^. The interaction interfaces between the subsequent subunits were found using the interfaces command within ChimeraX, which calculates the buried solvent-accessible surface area and displays the associated residues. The electrostatic surface was found by converting the PDB to a pqr file using PDB2PQR^91^. The pqr file contains the charges of each atom. This was then opened in ChimeraX and the internal electrostatic tool was used to calculate the surface. The hydrophobicity surface was also calculated using the internal tool.

### Statistical test and replicates

All statistical tests were two-sided Student’s t-tests adapted to assume that variances are not necessarily equal (i.e. Welch’s test) apart from one-sided F-tests used for analysis of the apparent diffusion coefficient. Replicates comprised at least 10 separate fields of view for each condition which generated reasonable estimates for the sampled stoichiometry and mobility distributions.

## Data availability

Experimental data used in this article can be accessed from the doi: 10.5281/zenodo.20559106. Docking data are at DOI: <u>10.15124/35e4e340-5aee-4fe0-b49b-eae2722a5914</u>. All plasmids are available upon request, subject to restrictions including completion of materials transfer agreements.

## Code availability

All code used for instrument design/control, data acquisition/processing/analysis and figure generation directly relied on: MATLAB, LabVIEW, ImageJ and Python, source code available at https://github.com/pol-biophysics under Creative Commons Attribution-NonCommercial-ShareAlike 4.0 International License (CC-BY-NC-SA; https://creativecommons.org/licenses/by-nc-sa/4.0/).

## Acknowledgements

Many thanks to Dr Ji-Eun Lee (University of Edinburgh) for preliminary bioimaging investigations.

