## Supplementary Information for "DNA gyrase in live bacteria forms liquid condensates through weak multivalent bonding of excess GyrB"

### Supplementary Tables

| Strain (Relevant genotype) | LB medium | | | Minimal medium | | |
|---|---|---|---|---|---|---|
| | Mean doubling time (min) | s.d (min) | s.e.m (min) | Mean doubling time (min) | s.d (min) | s.e.m. (min) |
| Wild type (WT) | 28.35 | 1.03 | 0.59 | 83.31 | 4.08 | 2.35 |
| *gyrA-mCherry-<kan>* | 26.79 | 2.87 | 1.66 | 83.00 | 17.23 | 9.95 |
| *mGFP-gyrB* | 29.78 | 3.00 | 1.73 | 111.28 | 17.35 | 10.02 |
| *gyrA-mCherry-<kan> mGFP-gyrB* | 35.68 | 1.61 | 0.93 | 130.3 | 14.43 | 8.33 |

**Supplementary Table 1. Doubling times of labelled strains.** Total of n = 3 replicates performed for each strain, OD measurements binned into 10 min interval time points, standard deviation (s.d.) and standard error of the mean (s.e.m.) indicated.

| | Untreated | | | Rifampicin | | | Coumermycin A1 | | | Ciprofloxacin | | |
|---|---|---|---|---|---|---|---|---|---|---|---|---|
| | Mean ±sem | Number foci | Number cells | Mean ±sem | Number foci | Number cells | Mean ±sem | Number foci | Number cells | Mean ±sem | Number foci | Number cells |
| GyrA stoichiometry (all foci) | 19.85 ±1.14 | 217 | 81 | 22.61 ±1.22 | 140 | 46 | 31.64 ±1.11 | 366 | 121 | 23.79 ±0.77 | 400 | 143 |
| GyrB stoichiometry (all foci) | 50.63 ±1.77 | 223 | 81 | 57.00 ±2.53 | 137 | 46 | 61.19 ±1.95 | 254 | 121 | 47.34 ±1.18 | 375 | 143 |
| GyrA stoichiometry (colocalised) | 20.77 ±1.45 | 140 | 81 | 24.76 ±1.52 | 88 | 46 | 35.58 ±1.63 | 190 | 121 | 25.61 ±1.04 | 231 | 143 |
| GyrB stoichiometry (colocalised) | 53.33 ±2.38 | 121 | 81 | 57.45 ±3.37 | 74 | 46 | 64.5 ±2.47 | 155 | 121 | 49.71 ±1.83 | 190 | 143 |
| GyrA stoichiometry (not colocalised) | 18.18 ±1.83 | 77 | 81 | 18.98 ±1.98 | 52 | 46 | 27.39 ±1.42 | 176 | 121 | 21.31 ±1.14 | 169 | 143 |
| GyrB stoichiometry (not colocalised) | 47.44 ±2.64 | 102 | 81 | 56.46 ±3.85 | 63 | 46 | 56.00 ±3.13 | 99 | 121 | 44.92 ±1.45 | 185 | 143 |
| GyrA diffusivity (all foci) | 0.50 ±0.05 | 217 | 81 | 0.54 ±0.06 | 140 | 46 | 0.29 ±0.03 | 366 | 121 | 0.34 ±0.03 | 400 | 143 |
| GyrB diffusivity (all foci) | 0.27 ±0.03 | 223 | 81 | 0.20 ±0.04 | 137 | 46 | 0.20 ±0.02 | 254 | 121 | 0.20 ±0.02 | 375 | 143 |
| GyrA diffusivity (colocalised) | 0.57 ±0.07 | 140 | 81 | 0.57 ±0.08 | 88 | 46 | 0.27 ±0.03 | 190 | 121 | 0.34 ±0.03 | 231 | 143 |
| GyrB diffusivity (colocalised) | 0.26 ±0.04 | 121 | 81 | 0.16 ±0.03 | 74 | 46 | 0.23 ±0.04 | 155 | 121 | 0.18 ±0.02 | 190 | 143 |
| GyrA diffusivity (not colocalised) | 0.38 ±0.07 | 77 | 81 | 0.50± 0.09 | 52 | 46 | 0.31 ±0.04 | 176 | 121 | 0.34 ±0.04 | 169 | 143 |
| GyrB diffusivity (not colocalised) | 0.29 ±0.06 | 102 | 81 | 0.25 ±0.07 | 63 | 46 | 0.14 ±0.03 | 99 | 121 | 0.21 ±0.03 | 185 | 143 |

**Supplementary Table 2. Mean and s.e.m. values for stoichiometry and diffusivity for GyrA and GyrB of all tracked clusters.**

| Condition | *a* | *Δa* | *b (/s)* | *Δb (/s)* | *c* | *Δc* | $t_{1/2}$ (s) | $\Delta t_{1/2}$ (s) |
|---|---|---|---|---|---|---|---|---|
| **Gyr A Untreated** | 0.408 | 0.058 | 2.265 | 0.821 | 0.419 | 0.034 | 0.306 | 0.111 |
| **GyrB Untreated** | 0.542 | 0.050 | 2.858 | 0.644 | 0.572 | 0.027 | 0.243 | 0.055 |
| **GyrA + HEX** | 0.254 | 0.033 | 0.997 | 0.362 | 0.251 | 0.025 | 0.695 | 0.253 |
| **GyrB + HEX** | 0.373 | 0.047 | 0.504 | 0.180 | 0.419 | 0.045 | 1.376 | 0.492 |

**Supplementary Table 3 Fitting outputs for a one-exponential FRAP model.** Normalised FRAP intensity $I$ values were fitted using a model which assumes an exponential recovery weighted by parameter *a* of time constant *b* (= ln2/$t_{1/2}$) with steady-state recovery level of *c* through the equation $I = c - aexp(-bt)$. Fitting error values are quoted at 95% confidence intervals (1 s.d.)

| Structure | | Subunits | | Docking Score | Confidence Score |
|---|---|---|---|---|---|
| | | GyrA | GyrB | | |
| GyrB + GyrB | | 0 | 2 | -240.45 | 0.859 |
| GyrA + GyrA | | 2 | 0 | -468.27 | 0.998 |
| GyrA +GyrB | | 1 | 1 | -575.87 | 0.9998 |
| **1** | 2(GyrA + GyrB) | 2 | 2 | -424.81 | 0.996 |
| **2** | **[1]** + GyrB | 2 | 3 | -232.64 | 0.839 |
| **3** | **[2]** + GyrB | 2 | 4 | -260.40 | 0.901 |
| **4** | **[3]** + 2(GyrA + GyrB) | 4 | 6 | -231.07 | 0.835 |
| **5** | **[4]** + GyrB | 4 | 7 | -237.64 | 0.852 |
| **6** | **[5]** + GyrB | 4 | 8 | -233.75 | 0.842 |

**Supplementary Table 4. Docked structures with a breakdown of how many of each subunit are in each structure as well as the docking and confidence scores.** Docking of the individual subunits and the building up of the cluster are shown, [number] represents the structure being added to. Colour scheme corresponds to that used in Fig. 5. Docking scores are unitless empirical values with more negative values indicating energetically more favourable states. The confidence score is a unitless P value such that values of 0.7 or higher indicate a heuristic threshold for likely binding.

| Structure | Docking Score | Confidence Score |
|---|---|---|
| GyrA + DNA | -311.05 | 0.962 |
| GyrB + DNA | -332.26 | 0.975 |
| 2(GyrA) + DNA | -290.14 | 0.943 |
| 2(GyrB) + DNA | -367.31 | 0.987 |
| 2(GyrA + GyrB) + DNA | -325.31 | 0.971 |

**Supplementary Table 5. Docking and confidence scores for individual subunits and homodimers docking to relaxed circular DNA.** More negative values for the (unitless) docking score indicate greater energetic favourability and higher (unitless) P values shown by the confidence score.

| Plasmid | Description | Antibiotic | Reference |
|---|---|---|---|
| pCP20 | Yeast Flp recominase expression plasmid | Amp | [1] |
| pKD46 | λ Red recombinase expression plasmid | Amp | [1] |
| pAP1 | pUC18 linker-mGFP-<kan> | Amp, kan | This study |
| pJGB374 | pUC18 linker-mCherry-<kan> | Amp, kan | [2] |

**Supplementary Table 6. Plasmids used in this study to generate fluorescently tagged strains.** References refer to the Supplementary References in Supplementary Material.

| Strain | Relevant genotype | Source or derivation |
|---|---|---|
| TB28 | *ΔlacIZYA* | [3,4] |
| AS97 | TB28/pKD46 | [2] |
| AS901 | *<kan>-mGFP-gyrB* | *<kan>-mGFP* with homology to *gyrB* recombineered into AS97 |
| AS905 | *<>-mGFP-gyrB* | AS901 to $km^{s}$ with pCP20 |
| AS909 | *gyrA-mCherry-<kan>* | *mCherry-<kan>* with homology to *gyrA* recombineered into AS97 |
| AS912 | *gyrA-mCherry-<kan> <>-mGFP-gyrB* | AS905 X P1(AS909) to $km^{R}$ |

**Supplementary Table 7. Strains used in this study.** All strains were derivatives of the common laboratory strain MG1655. References refer to the Supplementary References in Supplementary Material.

| Primer | Sequence (5’ - 3’) |
|---|---|
| oAS182 | ACGATGAAATCGCTCCGGAAGTGGACGTTGACGACGAGCCAGAAGAAGAAGGCTG GCTCCGCTGCTGG |
| oAS183 | TTTCAATTCAAACAAGGGAGATAGCTCCCTTTTGGCATGAAGAAGTAAAACATATGAAT ATCCTCCTTAG |
| oAS188 | TTATCTACCACCTCGAATAC |
| oAS195 | AAAGGGTAAAATAACGGATTAACCCAAGTATAAATGAGCGAGAAACGTTGGTGTAGG CTGGAGCTGCTTC |
| oAS196 | GCATCCAGCCCTTTCAGGACTTTGATACTGGAGGAGTCATAAGAATTCGAATGAATTC GCCAGAACCAG |
| oJL1 | CGCGGAAGGGGACGATGAAATC |
| oJL14 | CACTCATTGCCACATTCC |
| oJL15 | CTCTGAGCTTGATGATGAG |
| oJGB398 | TTCAAGTGGGAACGCGTAATG |
| oJGB399 | CATCGTCTTCTTCTGCATCAC |
| oJGB417 | TACAAGACACGTGCTGAAGTC |
| oJGB418 | TGCTAGTTGAACGCTTCCATC |
| oMKG70 | CAGTCATAGCCGAATAGCCT |
| oMKG71 | CGGTGCCCTGAATGAACTGC |
| oMKG72 | CGGCCACAGTCGATGAATCC |

**Supplementary Table 8. Primers used in this study.**

**Supplementary Figures**

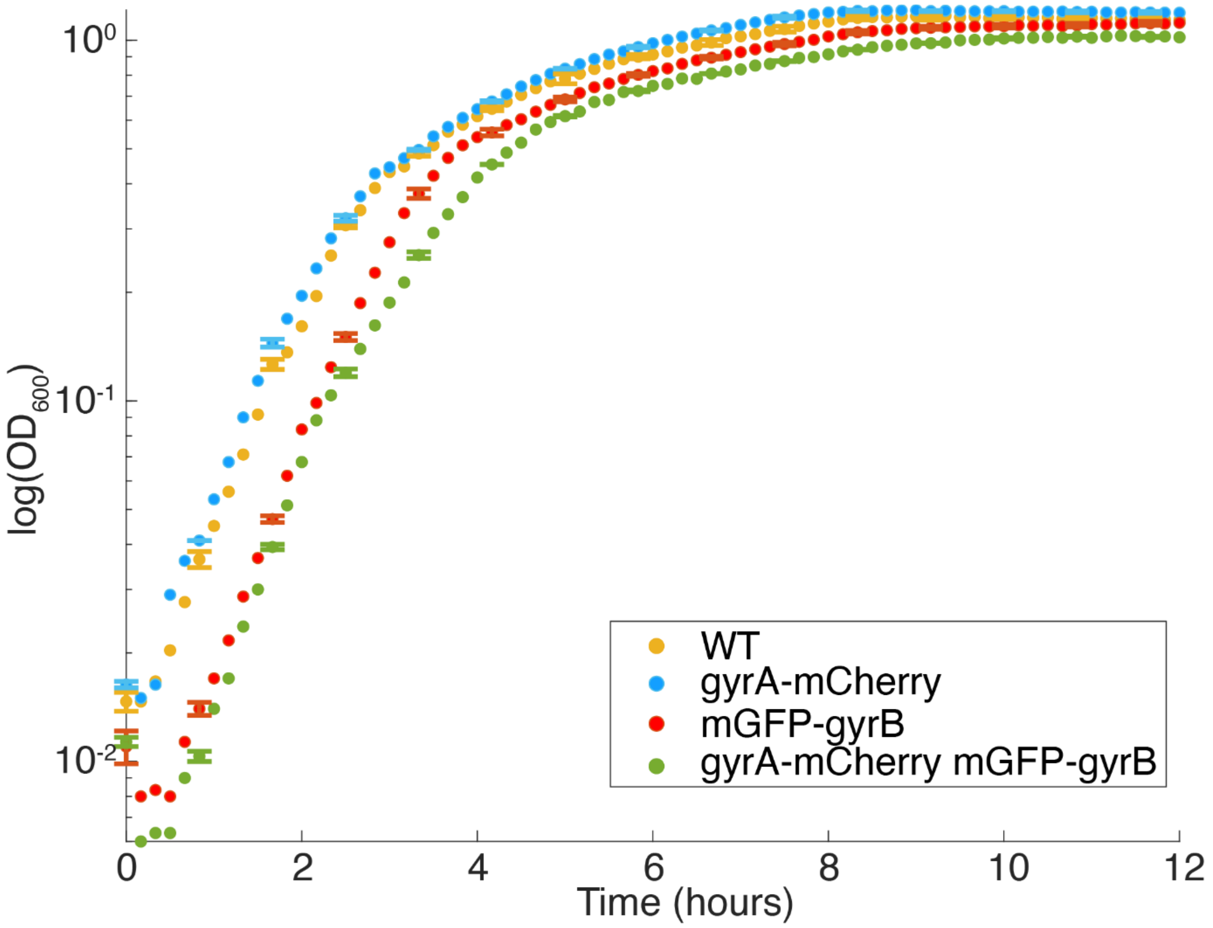


**Supplementary Figure 1. Growth curves in LB medium indicate only a marginal impairment to growth due to the presence of fluorescent tags.** Cultures were grown for 12 h in LB. $OD_{600}$ values are plotted on a log scale on the vertical axis with a linear scale of time on the horizontal axis. Relevant genotypes of the strains are indicated within the figure; s.d. error bars (shown just on every 5th consecutive point for clarity), taken from n = 3 replicate cultures.

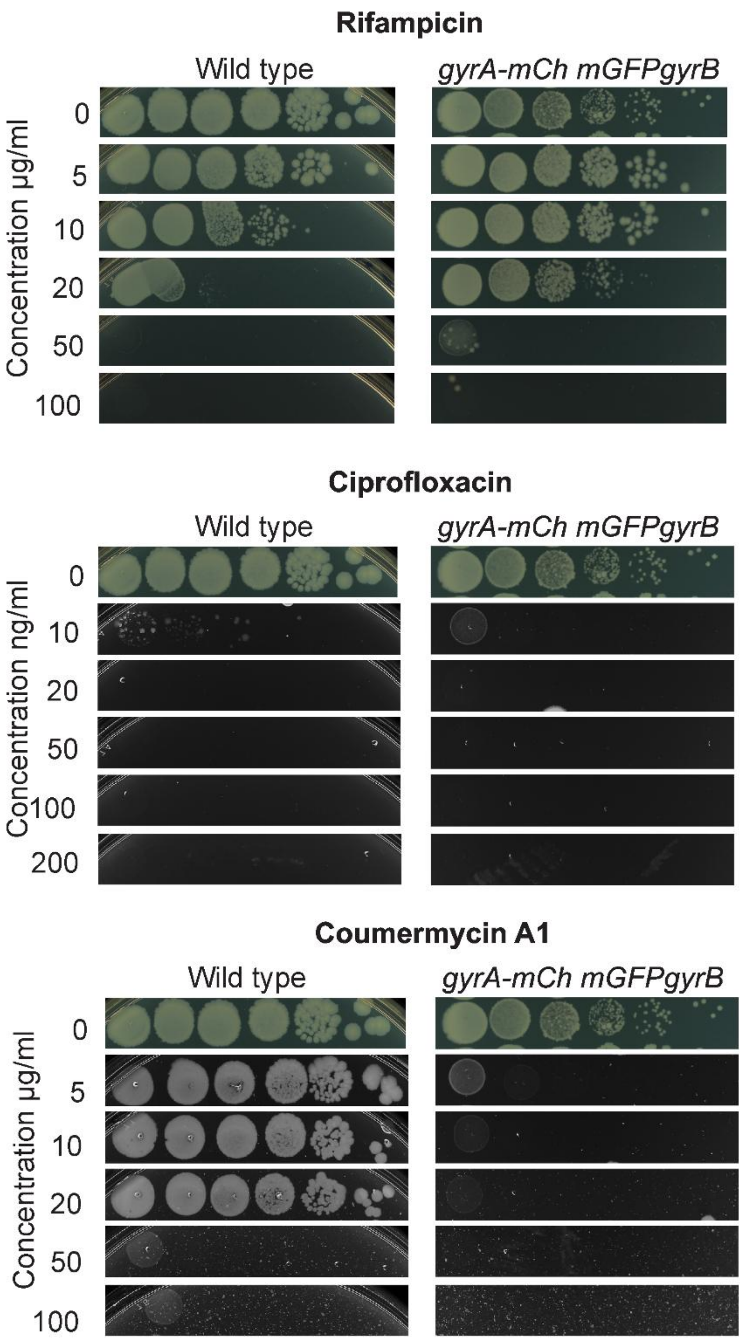


**Supplementary Figure 2. The antibiotic sensitivity of the GyrA-mCherry and mGFP-GyrB dual labelled strain compared with wild type.** Results of spot tests using the dual-labelled strain compared against the unlabelled isogenic wild type parent TB28 as a control, using LB plates containing antibiotics rifampicin, ciprofloxacin or coumermycin A1 at the concentrations indicated.

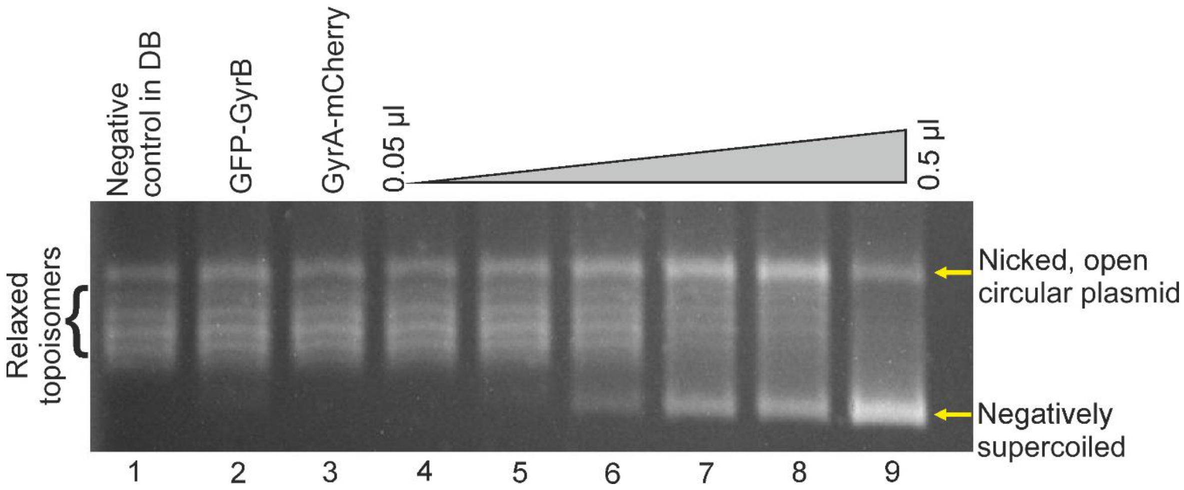


**Supplementary Figure 3. Supercoiling assays on purified fusion constructs indicate a clear ability to (−) supercoil relaxed plasmid DNA.** In this supercoiling assay, the substrate is relaxed plasmid pBR322 which is then (−) supercoiled by the presence of functional gyrase. The relaxed topoisomers of the plasmid can be separated by agarose gel electrophoresis. Lane 1 is a negative control lane in dilution buffer (DB) containing relaxed topoisomers of plasmid DNA pBR322 substrate used for all lanes, lanes 2 and 3 contain the GFP-GyrB and GyrA-mCherry purified fusion constructs alone respectively, lanes 4-9 have a mixture of GFP-GyrB and GyrA-mCherry comprising equal volumes of 0.05, 0.01, 0.02, 0.025, 0.033 and 0.5 µl respectively, showing the clear appearance of a band corresponding to negatively supercoiled pBR322.

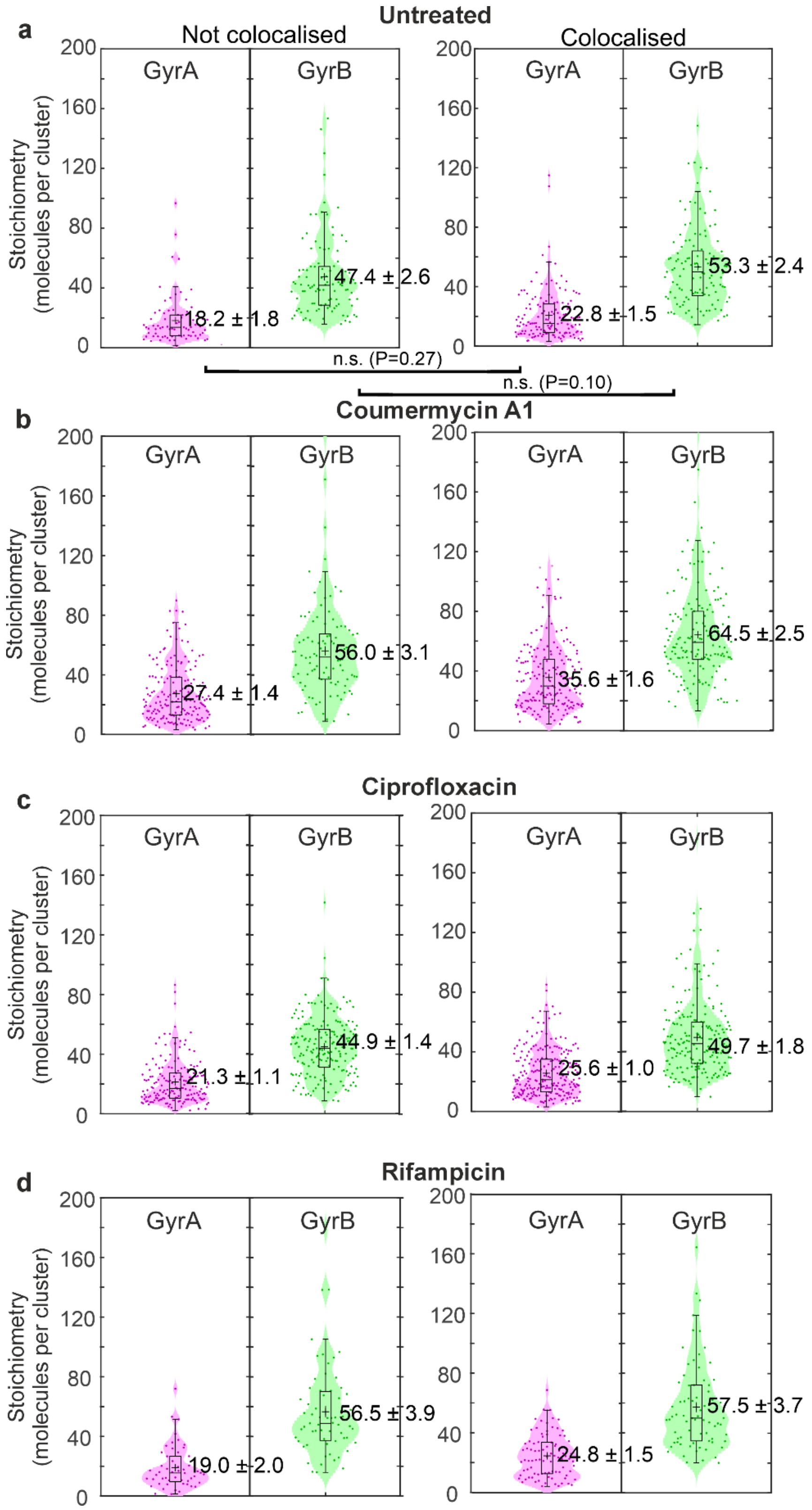


**Supplementary Figure 4. The relative abundance of GyrA and GyrB in clusters is largely insensitive to cluster colocalisation. a.** Violin plots showing stoichiometry distributions for GyrA and GyrB clusters that are not (left panel) and are (right panel) colocalised (from n = 217 GyrA and n = 223 GyrB clusters), mean ±s.e.m. indicated, with P values from corresponding Student's t-tests indicating no significant differences (n.s.) between corresponding GyrA and GyrB; number of cells n = 81. Equivalent violin plots for cell treated with **b.** coumermycin A1 (from n = 366 GyrA and n = 254 GyrB clusters, number of cells = 121), **c.** ciprofloxacin (from n =400 GyrA and n = 375 GyrB clusters, number of cells = 143), and **d.** rifampicin (from n = 140 GyrA and n = 137 GyrB clusters, number of cells = 46).

| DNA (pmol): | 1 | 1 | 1 | 1 | 1 | 1 |
|---|---|---|---|---|---|---|
| GyrA (pmol): | 0 | 2 | 4 | 2 | 4 | 10 |
| GyrB (pmol): | 0 | 10 | 20 | 2 | 4 | 10 |

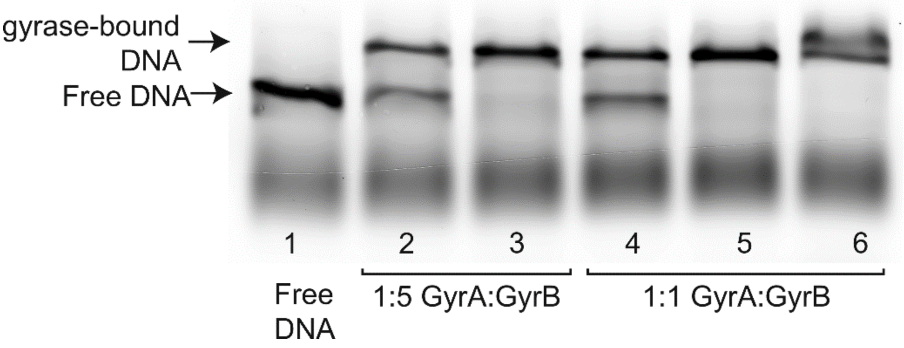


**Supplementary Figure 5. Excess GyrB does not significantly enhance DNA binding to gyrase *in vitro*.** Electrophoretic Mobility Shift Assays (EMSA) were performed using a Cy5-labelled 167 bp DNA fragment (1 pmol in each lane). For each sample (1-6), GyrA and GyrB were first pre-mixed at either canonical (1:1) or a 1:5 molar ratio, followed by incubation with DNA for 1 h at 37 C, and electrophoresed. At matched GyrA amounts, increasing GyrB 5-fold (lanes 2 vs 4, or 3 vs 5) did not lead to a measurable change of the proportion of gyrase-bound DNA to within ~5%.

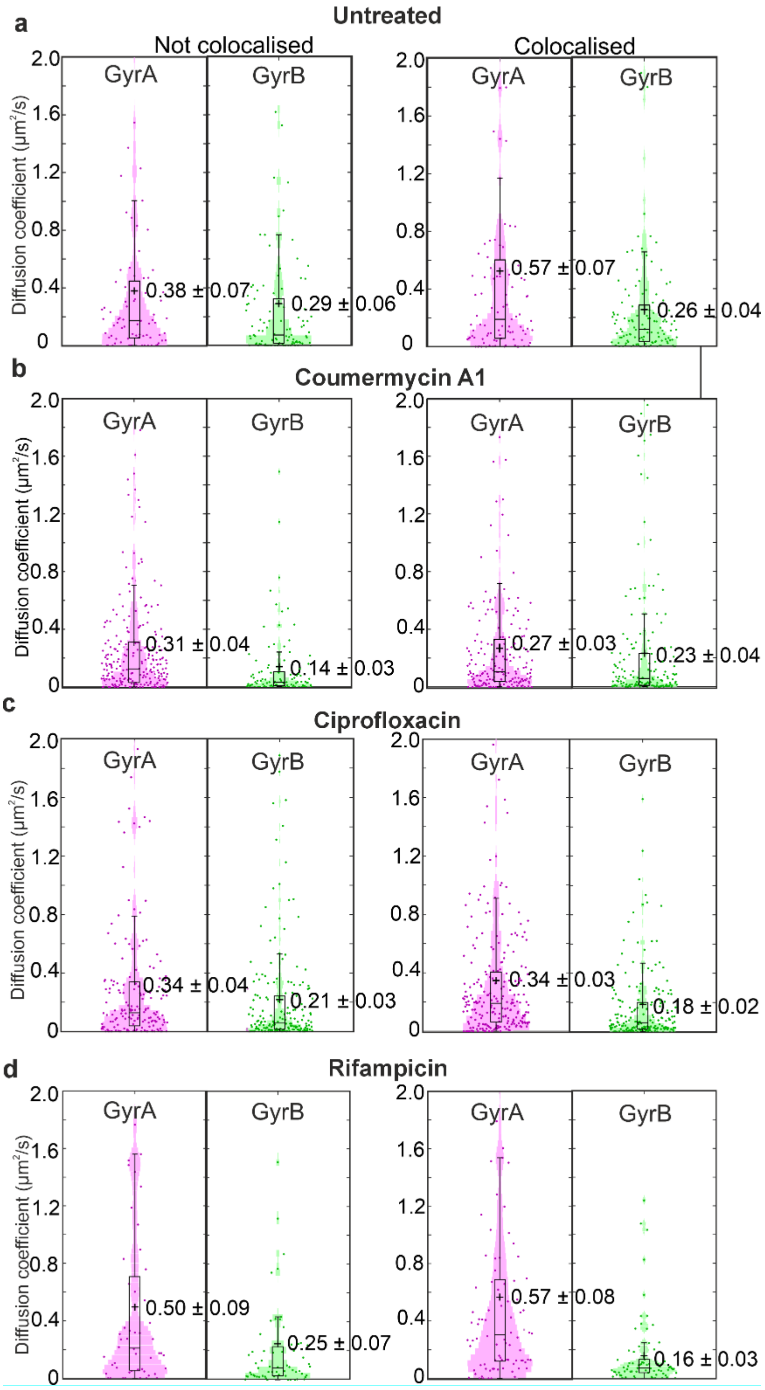


**Supplementary Figure 6. The mobility of GyrA clusters is higher than that of GyrB whether or not GyrA and GyrB are colocalised.** Violin plots with overlaid jitter data of diffusivity for the GyrA and GyrB components of clusters which are (right panels) (n = 140 GyrA clusters and 121 GyrB clusters) and are not (left panels) (n = 77 GyrA clusters and 102 GyrB clusters) colocalised. **b.** coumermycin A1 (n = 190 GyrA clusters, n = 155 GyrB clusters colocalised; n = 176 GyrA clusters, n = 99 GyrB clusters not colocalised), **c.** ciprofloxacin (n = 231 GyrA clusters, n = 190 GyrB clusters colocalized; n = 169 GyrA clusters, n = 185 GyrB clusters not colocalised) and **d.** (n = 88 GyrA clusters, n = 74 GyrB clusters colocalised; n = 52 GyrA clusters, n = 63 GyrB clusters not colocalised) treated cells; means (grey dash lines) ±s.e.m. indicated.

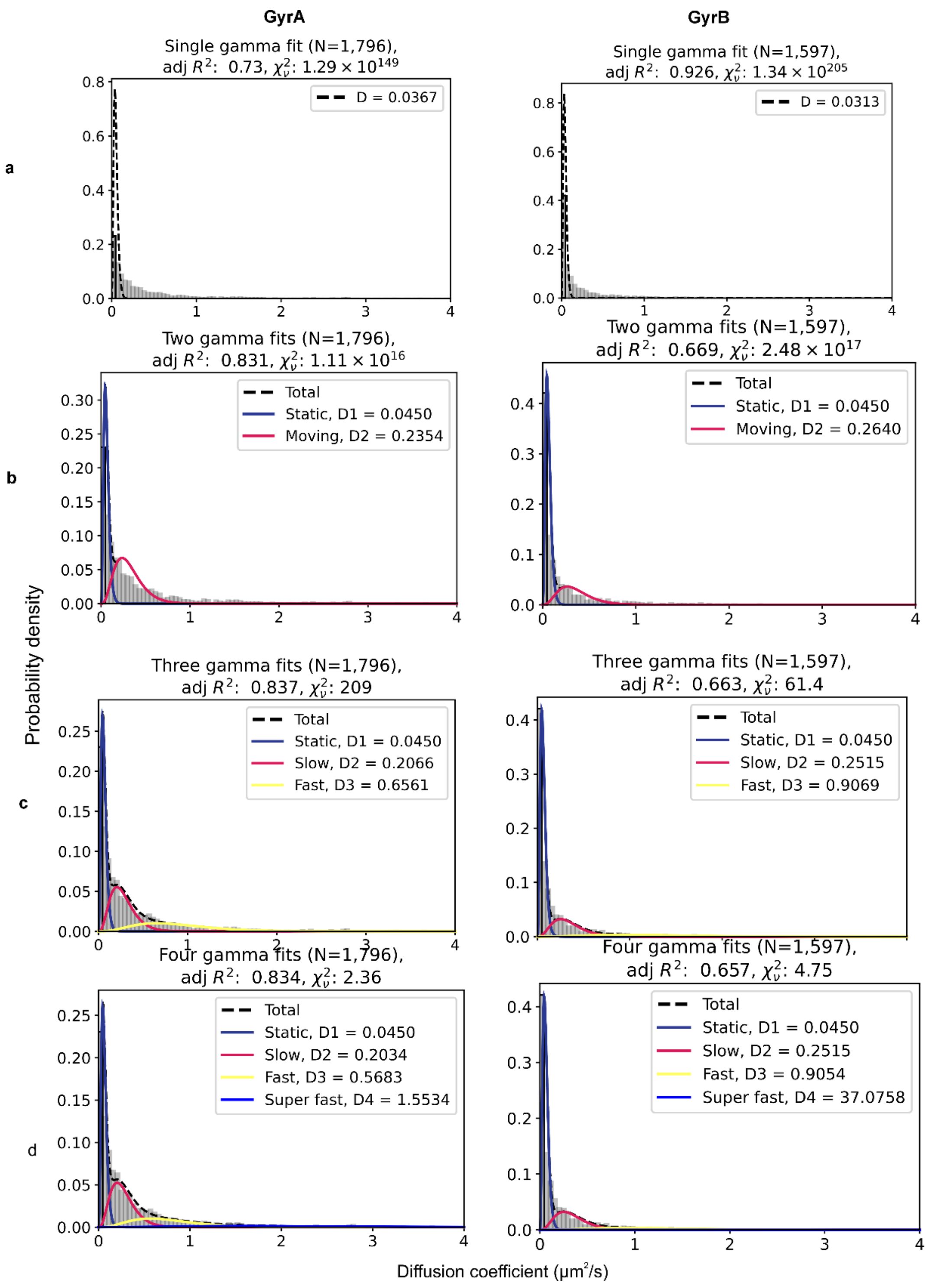


**Supplementary Figure 7. The distribution of diffusion coefficients for GyrA and GyrB clusters can be fitted using multiple mobility components**. Fits to combined data for the GyrA component (left column) and GyrB component (right column) of clusters indicated for **a.** 1-component, **b.** 2-component, **c.** 3-component and **d**. 4-component Brownian diffusion mode contributions, with the optimised modal values of apparent diffusion coefficient *D* indicated along with adjusted $R^2$ and reduced $\chi^2$ values for the total fits for each.

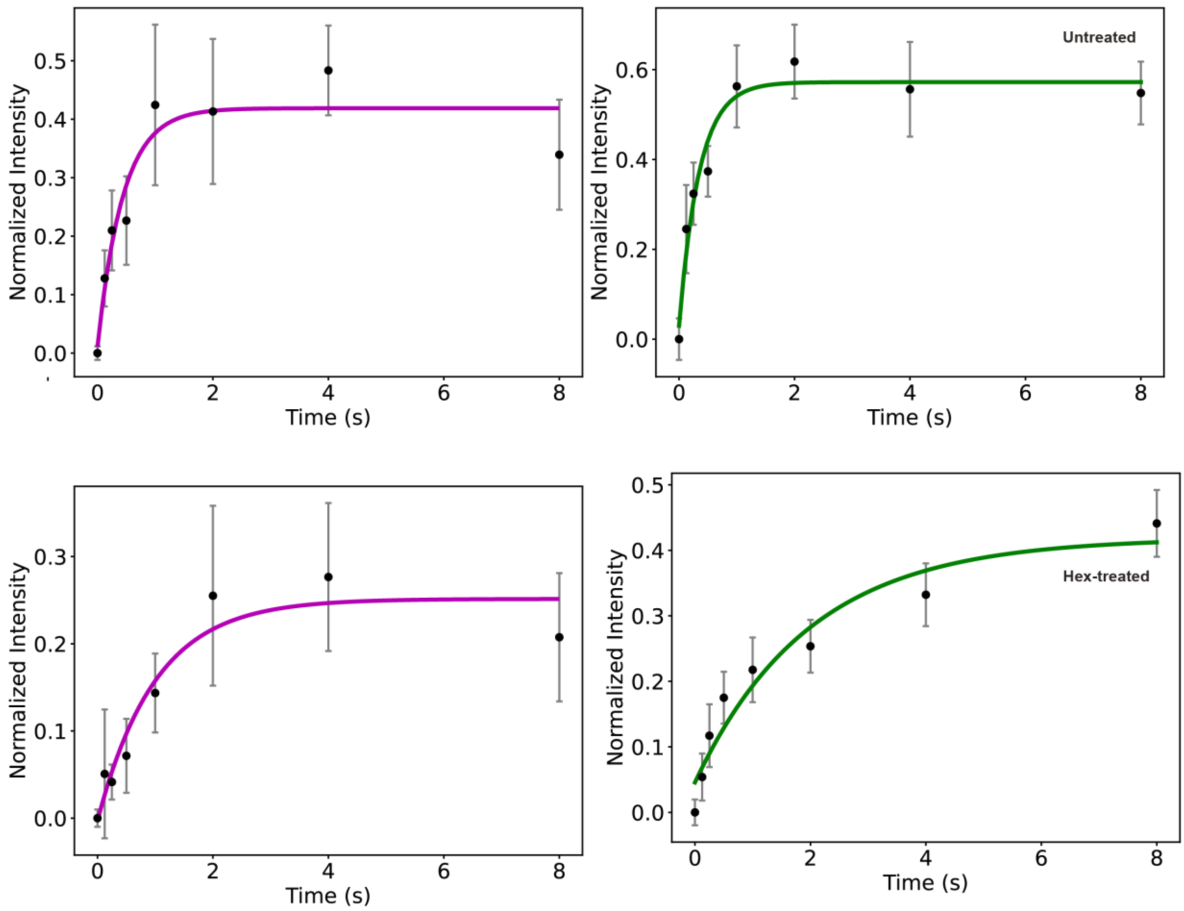


**Supplementary Figure 8.** FRAP recovery analysis using one-component exponential recovery fits for GyrA (magenta) and GyrB (green), s.e.m. error bars, for untreated (upper panels) HEX treated (lower panels) cells. Number of traces n = 20-24.

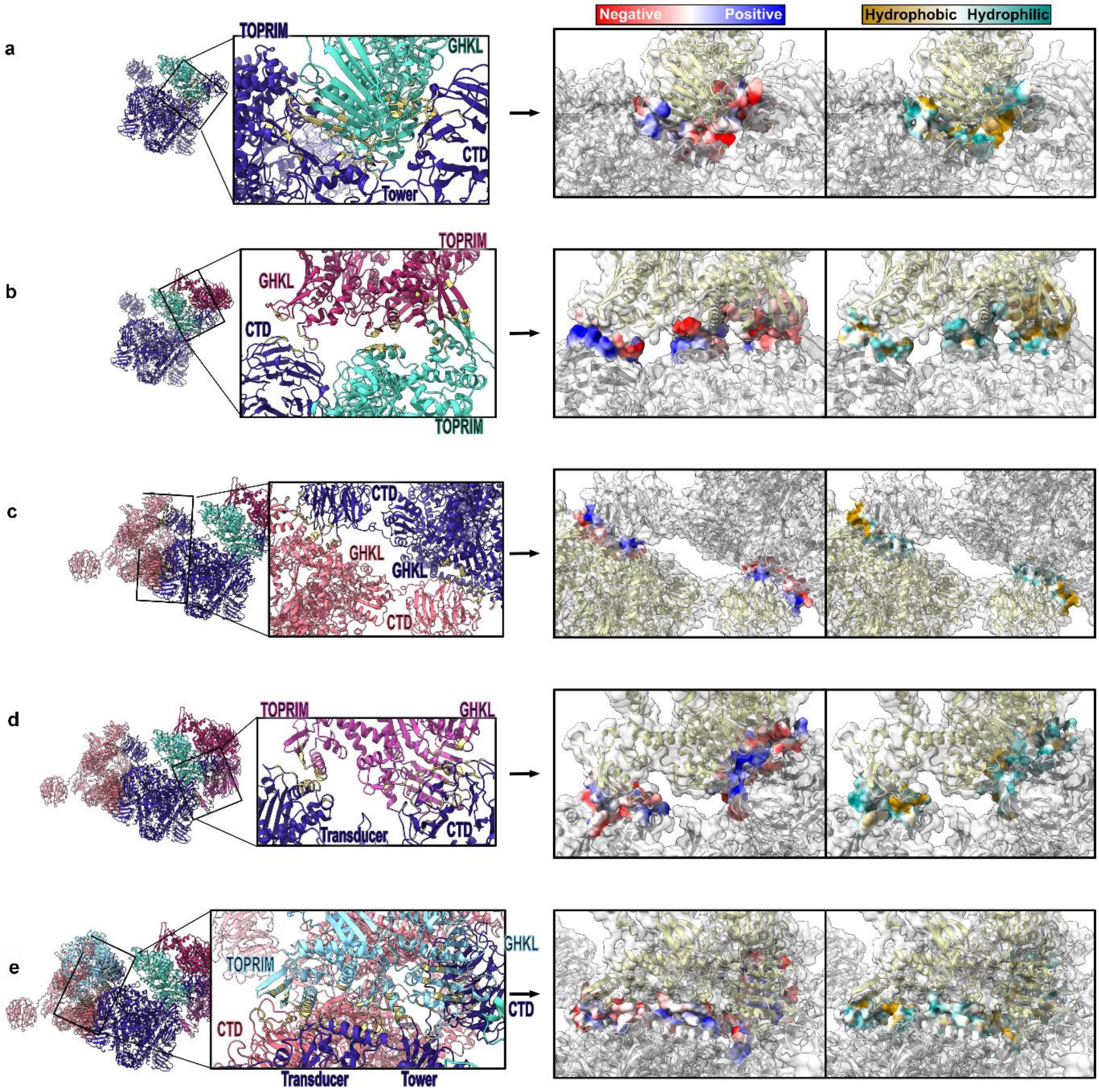


**Supplementary Figure 9. Excess GyrB forms weak, multivalent interactions between gyrase heterotetramers via several different residues.** Snapshots from corresponding Supplementary Movies 1-5 showing clustered structure following structural docking containing two heterotetramers and four individual GyrB subunits with an overall A:B stoichiometry ratio of 1:2. Left panels show each subunit/heterotetramer in a different colour- first tetramer in dark blue, first docked GyrB subunit in turquoise, second GyrB in dark pink, second tetramer in light pink, third GyrB in purple, and the fourth and final GyrB subunit in light blue, interacting residues between all regions indicated (yellow). Right panels show zoom-ins of interaction interfaces with overlaid electrostatic (red/blue) and hydrophobicity (cyan/yellow) potentials of the interacting residues. **a.** Interactions between heterotetramer and first added GyrB subunit; GyrB subunit's GHKL domain docks to the heterotetramer's GyrA CTD and Tower domains, alongside its GyrB TOPRIM domain. **b.** Interactions between second added GyrB subunit and existing structure of panel a; it interacts with the first heterotetramer and first added GyrB subunit, with the GHKL of the second subunit interacting with the CTD of the first heterotetramer, and the TOPRIM insert of the second subunit interacting with the TOPRIM insert of the first subunit. **c.** Interactions between second heterotetramer and existing structure of panel b. It interacts with first heterotetramer; the GyrA CTD of the second heterotetramer interacts with the GyrB GHKL of the first heterotetramer and vice versa. **d.** Interactions between third added GyrB subunit and the existing structure of panel c. It interacts with the first heterotetramer; the TOPRIM domain of the third added GyrB subunit interacts with the Transducer region of the first heterotetramer, and the GHKL of the third subunit interacts with the CTD of the first tetramer. **e.** Interactions between fourth and final added GyrB subunit and the existing structure. It interacts with both heterotetramers, docking in the joint of the interacting tetramers. The TOPRIM domain of the fourth individual subunit interacts with the Transducer of the first heterotetramer and the CTD of the second, and the GHKL of the subunit interacts with the CTD of the first heterotetramer.

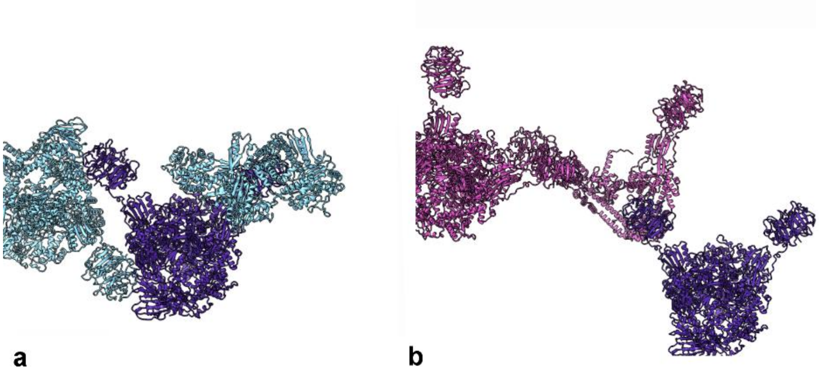

a b

**Supplementary Figure 10. Comparison of docking with excess GyrB vs. excess GyrA.** Panel **a** shows excess GyrB and is taking the first four steps of the cluster built up before - adding two GyrB subunits, then a second tetramer (A:B ratio = 2:3). The first tetramer is shown in dark blue, and the added GyrB subunits and tetramer are shown in light blue. They can be seen to form more of a ball-shaped cluster. Panel **b** shows the excess GyrA scenario (A:B = 3:2), where a GyrA homodimer was docked, then a second tetramer. These are both shown in pink, with the first tetramer again is shown in dark blue. They can be seen to form a more filament-like structure, with each new docking adding to the end of the previous.

**Supplementary Movies 1-5 legends**

The five videos (with associated screenshots) illustrate the stepwise docking process used in generating a gyrase cluster which ultimately comprises two heterotetramers and four individual GyrB monomer subunits with an overall A:B ratio of 1:2. Each subunit/tetramer is shown in a different colour- first tetramer in dark blue, first docked GyrB subunit in turquoise, second GyrB in dark pink, second tetramer in light pink, third GyrB in purple, and the fourth and final GyrB subunit in light blue. The interacting residues between all regions are shown in yellow. Each video contains a zoom in on the interaction which then has the hydrophobic surface fade in and out (yellow is hydrophobic, blue is hydrophilic), followed by the electrostatic surface (blue is positive, red is negative) before zooming out again.

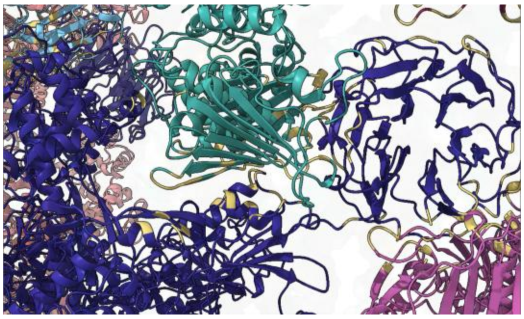

**Supplementary 1** Video shows an initial zoomed out view with the surfaces and a scale bar in the bottom left corner which then fades, revealing the cartoon structure. The video then zooms in to the first interaction between the heterotetramer and the first GyrB subunit, where the GyrB subunit's GHKL domain docks to the tetramer's GyrA CTD and Tower domains, alongside its GyrB TOPRIM domain. The surfaces are then shown, and the video zooms out again.

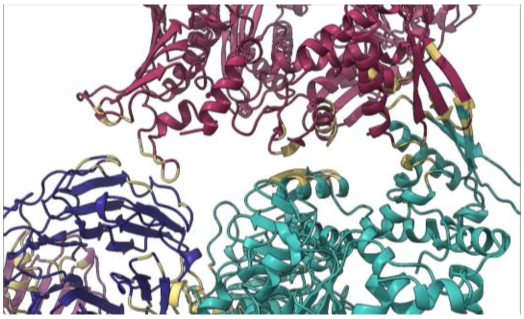

**Supplementary 2** Video shows the initial zoomed out cartoon structure which then zooms into the second interaction between the second GyrB subunit and the existing structure. It interacts with both the first heterotetramer and the first GyrB subunit, with the GHKL of the second subunit interacting with the CTD of the first tetramer, and the TOPRIM insert of the second subunit interacting with the TORPRIM insert of the first subunit. Both surfaces are shown, then the video zooms out again.

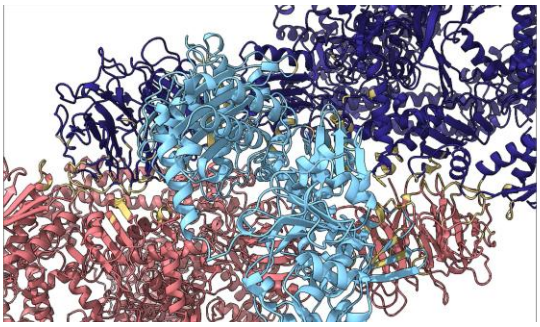

**Supplementary 3** Video shows the initial zoomed out cartoon structure which then zooms into the third interaction between the second tetramer and the existing structure. It interacts with the first heterotetramer. The GyrA CTD of the second tetramer interacts with the GyrB GHKL of the first tetramer and vice versa. Surfaces are shown and the video zooms back out.

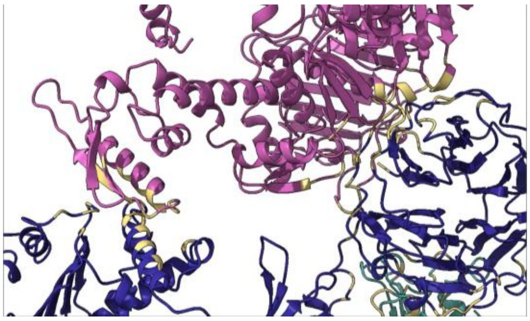

**Supplementary 4** Video shows the initial zoomed out cartoon structure which then zooms into the fourth interaction between the third GyrB subunit and the existing structure. It interacts with the first heterotetramer. The TOPRIM domain of the third GyrB subunit interacts with the Transducer region of the first tetramer, and the GHKL of the third subunit interacts with the CTD of the first tetramer. Both surfaces are shown, then the video zooms out again.

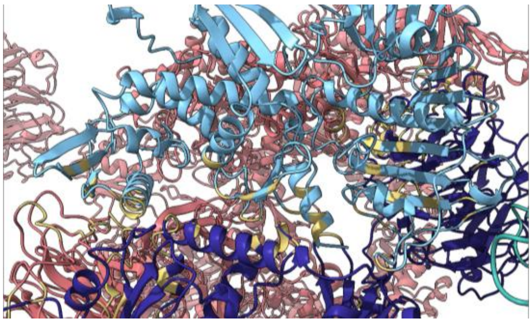

**Supplementary 5** Video shows the initial zoomed out cartoon structure which then zooms into the fifth interaction between the fourth and final GyrB subunit and the existing structure. It interacts with both heterotetramers, docking in the joint of the interacting tetramers. The TOPRIM domain of the fourth individual subunit interacts with the Transducer of the first tetramer and the CTD of the second, and the GHKL of the subunit interacts with the CTD of the first tetramer. Both surfaces are shown, then the video zooms out again and the overall surface and scale bar fade back in.
